# scRNA-seq-based analysis of skeletal muscle response to denervation reveals selective activation of muscle-resident glial cells and fibroblasts

**DOI:** 10.1101/2020.12.29.424762

**Authors:** C Nicoletti, X Wei, U. Etxaniz, D Proietti, L. Madaro, P.L. Puri

## Abstract

Developmental synaptogenesis toward formation of neuromuscular junctions (NMJs) is regulated by the reciprocal exchange of signals derived from nerve or muscle ends, respectively. These signals are re-deployed in adult life to repair NMJ lesions. The emerging heterogeneity of skeletal muscle cellular composition and the functional interplay between different muscle-resident cell types activated in response to homeostatic perturbations challenge the traditional notion that muscle-derived signals uniquely derive from myofibers. We have used single cell RNA sequencing (scRNA-seq) for a longitudinal analysis of gene expression profiles in cells isolated from skeletal muscles subjected to denervation by complete sciatic nerve transection. Our data show that, unlike muscle injury, which massively activates multiple muscle-resident cell types, denervation selectively induced the expansion of two cell types - muscle glial cells and activated fibroblasts. These cells were also identified as putative sources of muscle-derived signals implicated in NMJ repair and extracellular matrix (ECM) remodelling. Pseudo-time analysis of gene expression in muscle glial-derived cells at sequential timepoints post-denervation revealed an initial bifurcation into distinct processes related to either cellular de-differentiation and commitment to specialized cell types, such as Schwann cells, or ECM remodeling. However, at later time points muscle glial-derived cells appear to adopt a more uniform pattern of gene expression, dominated by a reduction of neurogenic signals. Consensual activation of pro-fibrotic and pro-atrophic genes from fibroblasts and other muscle-resident cell types suggests a global conversion of denervated muscles into an environment hostile for NMJ repair, while conductive for progressive development of fibrosis and myofiber atrophy.

## Introduction

Nerve-muscle interactions are mediated by reciprocal exchange of signals derived from the two essential components of the neuromuscular junctions (NMJs) – the terminal end of the motor neuron and the motor end plate of the innervated myofiber (Lepore et al., 2019). Nerve-derived and muscle-derived signals cooperate to guide nerve-to-muscle interactions for synaptogenesis and NMJ formation during development (Alvarez-Suarez et al., 2020; Barik et al., 2016; Darabid et al., 2014; Lin et al., 2001). Developmental signals are typically re-deployed in post-natal life to repair injured/damaged or diseased tissues (Snider and Tapscott, 2003). Indeed, during nerve repair a number of nerve-derived signals have been implicated in motor neuron/axon guidance for re-growth and regeneration (Rigoni and Negro, 2020). Nerve repair within the context of NMJ loss also relies on muscle-derived signals; however, the cellular source of these signals has not been yet clearly defined.

Irreversible denervation by sciatic nerve transection provides an optimal experimental setting, whereby interruption of motor neuron-derived anterograde signals is instrumental to separate muscle-derived from motor neuron-derived signals. Within this setting, longitudinal single cell-based transcriptional analysis of muscles post-denervation enables capturing cell type-specific transcriptional signatures that can identify denervation-activated muscle-resident cell types and predict their contribution to muscle-derived signals implicated in muscle response to denervation.

Mounting evidence indicates that skeletal muscle response to homeostatic perturbations involves the coordinated activation of a multitude of muscle-resident cell types that cooperate to restore the original homeostasis (Farup et al., 2015). Recent single cell-based transcriptional analyses have revealed the identity of muscle-resident cells and their dynamic heterogeneity in response to acute or chronic perturbations of skeletal muscle homeostasis (De Micheli et al., 2020; Dell’Orso et al., 2019; Giordani et al., 2019; Malecova et al., 2018; Oprescu et al., 2020; Pawlikowski et al., 2019; Petrany et al., 2020). These studies have established that the response to the most investigated type of homeostatic perturbation of skeletal muscles – the physical injury by myotrauma – is highly influenced by the inflammatory infiltrate, which transiently provides regulatory signals to other muscle-resident cells, including fibro-adipogenic progenitors (FAPs) and other interstitial cell types, to ultimately promote muscle stem cells (MuSCs)-mediated regeneration (Biferali et al., 2019; Farup et al., 2015). Lack of complete resolution of reactive inflammation and/or dysregulated interactions between inflammatory cells and muscle-resident cells have been implicated in the pathogenesis of many muscular diseases, including muscular dystrophies, in which chronic injury occurs as consequence of genetic mutations leading to repeated cycles of myofiber degeneration/regeneration following contraction (Judson et al., 2013; Malecova et al., 2018; Morgan and Partridge, 2020; Serrano and Muñoz-Cánoves, 2017).

By contrast, perturbation of muscle homeostasis by denervation is accompanied by a milder inflammatory response, activation of FAP-derived pro-atrophic signals and negligible activation of MuSCs-mediated regeneration (Madaro et al., 2018). Moreover, previous studies have implicated different cell types, including Schwann cells and MuSCs, in muscle responses to nerve injury, such as NMJ repair and ECM remodeling (Kang et al., 2003; Liu et al., 2015b, 2017). However, a systematic, unbiased identification of the muscle-resident cell types activated by denervation, and the activation of related functional networks, has not been performed to date. This gap of knowledge has significantly prevented our understanding of the pathogenesis of denervation-based muscular diseases and the identification of targets for therapeutic interventions toward repairing diseased or injured neuromuscular junctions.

Here we present a longitudinal analysis of the transcriptional profiles of muscle-resident cells at the single cell level (single cell RNA sequencing – scRNA-seq), upon acute denervation by sciatic nerve transection. This experimental strategy was instrumental to identify cell types and gene expression profiles activated upon interruption of motor neuron-derived signals, and enabled capturing putative cellular source of muscle-derived retrograde signals induced by denervation.

## Results

### scRNA-seq-mediated identification of denervation-responsive muscle-resident cell types

We used scRNA-seq analysis of whole skeletal muscles subjected to denervation, as compared to unperturbed counterparts, in order to capture gene expression signatures that could identify specific muscle-resident cells activated in response to denervation. We performed scRNA-seq of single cell suspensions prepared from 2 muscles downstream to sciatic nerve – tibialis anterior (TA) and gastrocnemius (GA) muscles – isolated from 3 month-old male mice, either unperturbed or subjected to denervation by complete sciatic nerve transection (Fig. 1A - top). We selected 3 timepoints post-denervation (p.d.), based on previous works reporting on the activation kinetics of muscle-resident cells by the first 48 hours post-denervation (day 2 p.d.) and the expansion of FAPs with persistent activation of STAT3-IL6 pathway at later time points (days 5 and 15 p.d.) that eventually leads to myofiber atrophy and fibrosis (Madaro et al., 2018). We generated scRNA-seq datasets from two biological replicates of a pool of TA and GA muscles isolated at our specific timepoints post-denervation (abbreviated as DEN2d, DEN5d and Den15d in figures and tables) or from unperturbed mice (abbreviated as Non-DEN in figures and tables). Fluorescence Activated Cell Sorting (FACS) was used to isolate single cells and eliminate contaminating cell debris, doublets and dead cells (Fig. S1A-D). Single cell libraries were generated with 10X Genomics Chromium Platform and sequenced on the Illumina Nova-Seq platform. We isolated a total of 48,666 cells across all the experimental conditions, with 44,403 cells analysed post-filtering, and detected a total of 36,185 genes (average of 6,000 cells per samples) with a read depth between 30k and 50k per sample (Fig. S1A-D). Seurat package (Stuart et al., 2019) was used for most of the data analysis. Data were filtered to exclude cells with higher rate of mitochondrial genes and cell doublets, as predicted with Scrublet (Wolock et al., 2019) (Fig. S1A-D). Principal Component Analysis (PCA), used for dimensionality reduction, revealed no batch effect between biological replicate samples from the different experimental points (Fig. S1E and G) or for cells captured at different cell cycle phases (Fig. S1F and H). Nevertheless, cells from all experimental points were normalized together taking into consideration their mitochondrial gene content and cell cycle-related genes.

**Figure 1.**
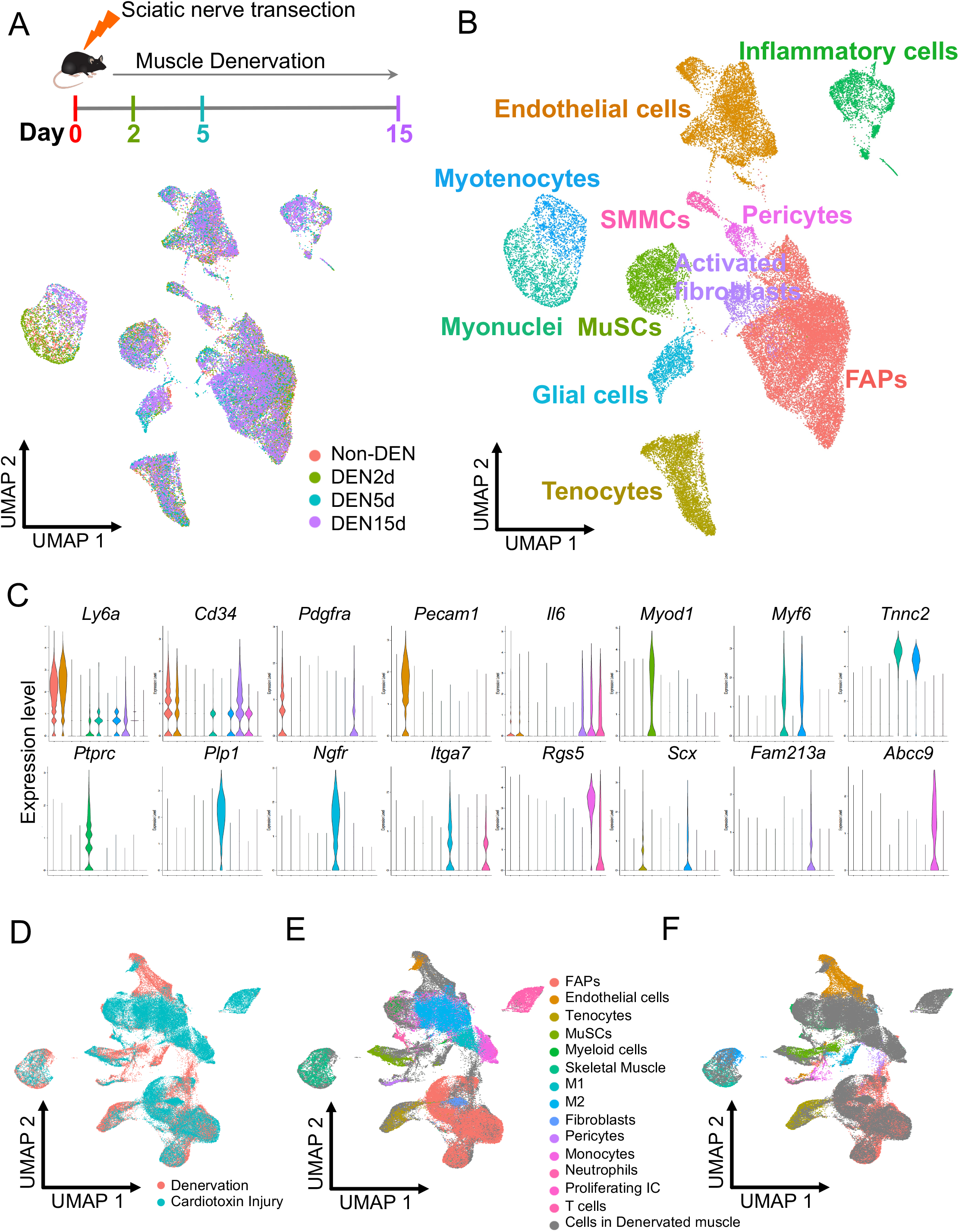
(A) Experimental Design. Single-cell suspensions were generated from whole TA and GA muscles from non-denervated mice and for three time points following denervation through sciatic nerve transection (namely, 0, 2, 5 or 15 days post-denervation, abbreviated as Non-DEN, DEN2d, DEN5d and Den15d in figures and tables). Cells from 3-months old male mice were isolated according to Rando’s protocol (Liu et al., 2015a), followed by FACS-sorting to remove cell debris, doublets and dead cells. UMAP embedding of the single-cell RNA-seq data colored by time point shows that changes in cell identity are restricted to few cell types directly involved in the denervation process. (B) UMAP embedding of all the single cells comprising the dataset colored by meta-cluster. (C) Violin plots of the expression profile of marker genes for each meta-cluster, color legend as in 1B. (D-F) UMAP embedding of the integration analysis of denervation and injury (Oprescu et al., 2020) datasets, with Oprescu et al. meta-cluster identities (E) or with the meta-cluster identities from our dataset (F-color legend as in 1B).

Independent unsupervised shared nearest neighbor (SNN) clustering was used to define clusters representing specific cell types and sub-types, and Universal Mani-fold Approximation and Projection (UMAP) was used as non-linear dimensionality reduction method to represent single cell transcriptomes from all conditions in the 2D space. These analyses identified 18 clusters of single cells, likely representing cellular states of muscle-resident cell types isolated from unperturbed muscles or at defined time points following denervation (Fig. 1A - bottom). Marker identification analysis found cell-type specific gene expression patterns, which revealed the expression of well-known lineage markers and further reduced single cell distribution into 11 meta-clusters (Fig. 1B). This analysis revealed the identity of FAPs (based on the simultaneous expression of *Ly6a, Cd34 and Pdgfra* (Joe et al., 2010; Uezumi et al., 2010), as the most represented population of muscle-resident cells (Fig. 1B and C; Fig. S2). Other major clusters identified additional and well-known muscle-resident cell types, including endothelial cells (based on *Pecam1* and *Ly6a* co-expression) (Albelda et al., 1991), pericytes (marked by the co-expression of *Abcc9, Pdgfrb* and *Kcnj8*) (Vanlandewijck et al., 2018), inflammatory cells (a heterogenous group of immune cells that share the selective expression of *Ptprc* - CD45) (Hermiston et al., 2003) and tenocytes (enriched for the expression of *Scx* and *Pdgfrb*) (Murchison et al., 2007) (Fig. 1B and C; Fig. S2). MuSCs were identified as cluster of heterogeneous cells expressing *Pax7, Myod1* and *Calcr* in various combinations (Baghdadi et al., 2018; Fukada et al., 2007; Seale et al., 2000) (Fig. 1B and C; Fig. S2). We also detected a cluster of cells that were identified as myonuclei, likely derived from a contamination of our cell preparation. They clustered adjacent to myotenocytes, as they both shared the selective expression of *Tnnc2*, but differ on the expression of *Scx*, which was restricted to myotenocytes (Fig. 1B and C; Fig. S2). Lastly, three additional clusters were assigned to recently identified muscle-resident cell types, such as smooth muscle and mesenchymal cells (SMMCs), activated fibroblasts and muscle glial cells (Fig. 1B and C; Fig. S2) (De Micheli et al., 2020; Giordani et al., 2019). For single cells dispersed between clusters, we could not identify any cell-type specific transcriptional signature or identity marker that could discriminate them from other muscle-resident cells. As these clusters share gene expression with surrounding cells types, we speculate that they could be composed by cells in dynamic transition from one cluster to another, within the spectrum of adjacent cell lineages.

Integrative analysis of our denervation dataset with a publicly available scRNA-seq dataset of muscle regeneration after cardiotoxin injury (Oprescu et al., 2020) demonstrates a large overlap between the muscle-resident cell types identified following denervation and those observed in previous studies on skeletal muscles undergoing regeneration post-injury (Fig. 1D-F). Of note, unlike regenerating muscles, which exhibit extensive changes in the size of multiple cell clusters following myotrauma (De Micheli et al., 2020; Dell’Orso et al., 2019; Oprescu et al., 2020; Pawlikowski et al., 2019; Petrany et al., 2020), denervation did not appear to significantly alter the average number of cells in most of the clusters (S. Table 1). In this regard, we reasoned that scRNA-seq provides a tool to measure expansions or contractions of specific cell types, when performed at sequential timepoints following an experimental perturbation. Indeed, the relative percentage of cluster representation is a rough reflection of their relative amount within the whole number of single cells isolated, assuming that there is no bias in cell type-specific extraction preference. With the exception of myonuclei, whose extraction is clearly biased by their anatomical position inside fibers, muscle-resident cell extraction are unlikely to be biased toward any particular cell types, as also indicated by recent studies of whole muscles showing that scRNA-seq analysis could capture quantitative dynamics of muscle-resident cell expansion post-injury that were previously detected based on immunohistochemistry or FACS analyses (De Micheli et al., 2020; Dell’Orso et al., 2019; Oprescu et al., 2020; Pawlikowski et al., 2019; Petrany et al., 2020). For instance, a dramatic expansion of the inflammatory infiltrate and immune-resident cells occurs in the 48 hours immediately after myotrauma, with the inflammatory cells accounting for about 90% of the single cells (Oprescu et al., 2020). We argue that this massive expansion of one cell type invariably reduces the capture of other muscle-resident cell types, which are by default underrepresented within the 10% of the non-inflammatory cell types analysed. Figure 2A and S. Table 1 show that unlike muscle injury by myotrauma, denervation did not trigger an immediate and massive infiltration of immune cells, which only by day 5 p.d. displayed a modest increase in number that remained unaltered by day 15 p.d. As such, denervated muscles did not exhibit any specific bias in the simultaneous analysis of muscle-resident cell types at any time point p.d. Another potential bias in the quantitative interpretation of longitudinal analysis of perturbed muscles by scRNA-seq could be provided by the massive loss of one or more cell types that account for a large proportion of cells, thereby skewing the representation of the other cell types (Fig. 2A and S. Table 1). Upon denervation, we observed the expected drastic reduction only of the myonuclei, whose number dramatically drops by day 5 p.d., after a paradoxical expansion observed at day 2 p.d. However, we note that myonuclei represent an accidental contamination from myofiber leakage typically occurring during single cell extraction, as also observed by others (De Micheli et al., 2020; Dell’Orso et al., 2019; Giordani et al., 2019; Oprescu et al., 2020). Since myonuclei account for less than 5% of cells post-filtering, they do not provide a significant bias. The transient drop of endothelial cell number observed at day 2 p.d., which coincided with a consensual increase in the proportion of FAPs, also did not significantly alter the percentage of cell type distribution (Fig. 2A; see also Fig. S3A).

**Figure 2.**
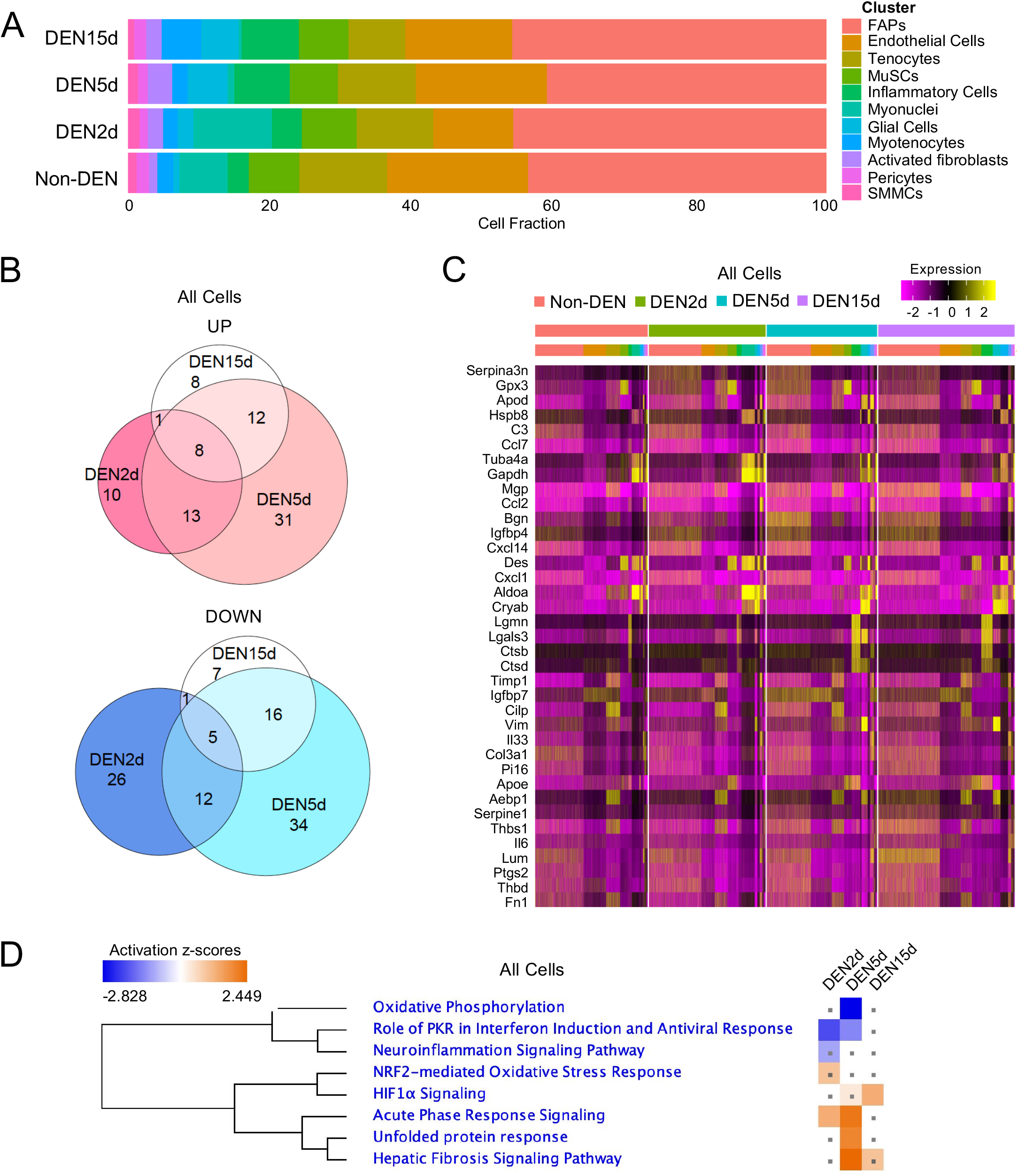
A) Barplot representing the proportion of single cells belonging to each meta-cluster for each experimental condition. (B) Venn diagram of upregulated genes (Upper) and downregulated genes (Bottom) in all single cells at DEN2d, DEN5d, and DEN15d compared with Non-DEN. DE Genes with p-value-adj <= 0.01 are counted. (C) Heatmap of top 20 upregulated genes in all cells at DEN2d, DEN5d, and DEN15d respectively compared with Non-DEN, clustered by condition and meta-cluster identity. (D) IPA comparison of DE genes in all cells at DEN2d, DEN5d, and DEN15d respectively compared with Non-DEN. Pathways significantly enriched in either time point with absolute z score > 1 were shown. Pathways with p-value >= 0.05 were indicated with gray dots.

Our scRNA-seq analysis revealed that, following denervation the majority of the largest and most represented clusters of muscle-resident cells did not exhibit dramatic alterations in their number; however, two cell types that were underrepresented in unperturbed muscles exhibited a progressively increase in both absolute number and percentage relative to all muscle-resident cells (Fig. 2A and S. Table 1). These putative denervation-activated populations coincided with clusters identified as muscle-resident glial cells and activated fibroblasts (Fig. 2A and S. Table 1). Their patterns of expansion were similar during the first 5 days p.d., with a progressive increase in both percentage and absolute cell number; however, by day 15 p.d. both cell types declined in term of percentage, despite the observed increase in the absolute number of glial cells (Fig. 2A and S Table 1). Progressive and moderate expansion of two cell types that are abundant in unperturbed muscles - myotenocytes and inflammatory cells – was also observed (Fig. 2A and S. Table 1). Modest fluctuations in FAPs and endothelial cells, tenocytes and SMMCs, were observed during our time-course post-denervation, while negligible changes in MuSCs number were observed at every time point post-denervation (Fig. 2A and S. Table 1).

The dynamic changes in the abundance of each cell type at sequential time points post-denervation is represented by the UMAP embedding shown in Figure S3A, which illustrates the selective and progressive expansion of muscle glial-derived cells and activated fibroblasts in response to denervation (Fig. S3A). At the same time, it shows that within other cell types that did not undergo significant expansions or reductions in number there were changes in internal distribution of gene expression, which predict transitions between cell states within the same lineage. Indeed, we found that sub-clusters identified within FAPs, endothelial cells and inflammatory cells underwent extensive reconfiguration along with the muscle response to denervation (Fig. S3A). We also determined the magnitude of transcriptional changes induced by denervation in each cell types, by comparing the number of differentially expressed (DE) genes in all cell types at each experimental condition. Supplementary Table 2 shows that the number of DE genes was higher in muscle glial-derived cells and activated fibroblasts (S. Table 2).

Overall, among all the muscle-resident cells, glial cells and activated fibroblasts were the cell types that exhibited the most significant changes in abundance and DE genes, indicating that they are the two major denervation-activated muscle resident cell types.

### Global dynamics of transcriptional responses of muscle-resident cells to denervation

Global gene expression analysis revealed that the magnitude of changes in gene expression of muscle-resident cells in response to denervation was much reduced when compared to the response to myotrauma. Venn diagram shows that the largest changes in gene expression occurred between days 2 and 5 p.d., while day 15 p.d. shows a reduced extent of DE genes (Fig. 2B). Functional analysis by IPA of DE genes across all the conditions predicted general trends of repression of oxidative phosphorylation, PKR-mediated viral response and neuroinflammation within the initial stages of muscle response to denervation – from days 2 to 5 p.d. - followed by activation of processes related to fibrosis and neo-angiogenesis that were detected at later stages – days 5 to 15 p.d. (Fig. 2C and D).

While these data indicate a general bi-phasic response of skeletal muscles to denervation, eventually leading to a persistent activation of fibrosis-related processes, global analysis of gene expression could be largely biased by the relative amounts of different cell types analyzed. In particular, a global analysis might not be informative on genes activated by denervation in cell types representing a very small percentage of the total muscle-resident cells, such as muscle glial cells and fibroblasts. Indeed, when we identified top marker genes for each cell type across all conditions (S. Table 3), we found that top genes of the largest populations were detected in the heatmap shown in Fig. 2C. For instance, top DE genes were enriched in ECM and inflammatory genes, which mostly derived from larger populations, such as FAPs, endothelial/pericytes and inflammatory cells. Conversely, top marker genes of the denervation-activated populations (muscle glial cells and activated fibroblasts), which represent a small percentage of the whole population of muscle-resident cells, were largely under-represented in the same heatmap (Fig. 2C and S. Table 3). Supplementary Figure 3B illustrates the relative contribution of multiple cell types, by day 5 p.d., to the expression levels of upregulated genes implicated in the major biological processes activated by denervation, such as ECM remodeling and fibrosis (Fig. S3B).

This prompted an interest to further perform functional analysis of differentially expressed genes by each individual muscle-resident cell type, in response to denervation.

### Muscle-resident cell type-specific responses to denervation

We have analysed the dynamic gene expression profiles of individual muscle-resident cell type identified by our scRNA-seq in response to denervation. Below are the analysis of individual cell types that were ranked based on their magnitude of activation (as defined by the changes in cell number and DE genes).

#### Muscle glial cells

Muscle glial cells are one of the least represented cell clusters in unperturbed muscles (Fig. 2A; Fig. S3A and S. Table 1); hence, their response to perturbations that trigger massive activation of larger populations (i.e. myotrauma) could not be entirely appreciated by previous scRNA-seq studies (De Micheli et al., 2020; Oprescu et al., 2020; Pawlikowski et al., 2019). Our analysis shows that muscle glial cells quickly responded to denervation with a progressive increase in cell number (Fig. 2A, Fig. S3A and S. Table 1) that was accompanied by significant changes in gene expression along with the time points p.d. (S. Table 2). Interestingly, most of DE genes were detected at day 2 and day 5 p.d., with a strong bias toward upregulated genes (Fig. 3A). Figure 3B displays the heatmaps of top DE genes at each experimental time point, showing a general upregulation of genes at days 2 and 5 p.d., followed by a consensual downregulation at day 15 p.d., although some genes, like *Igfbp3, Tnc, Vim* and *Btc*, exhibited a progressive increase from day 2 to day 15 p.d. (Fig. 3B). The glial identity of these cells was further confirmed by a comparative analysis with the Schwann cells markers used in earlier scRNA-seq dataset for glial cell-specific gene expression (Pawlikowski et al., 2019). Supplementary Fig. 4 shows that top Schwann cell type-specific genes extrapolated from this dataset were mostly enriched in our muscle-derived glial cells (Fig. S4). Furthermore, IPA analysis of DE genes across all experimental conditions predicted an activation of multiple neurogenic processes, including glial cell-derived neurotrophic factor (GDNF)- and Nerve Growth Factor (NGF)-mediated signaling, neuregulin signaling and other processes implicated in nerve repair, such as neuronal survival, synaptogenesis and axon guidance, between days 2 and 5 p.d. (Fig. 3C; S. Table 4) (Birchmeier and Bennett, 2016). However, these neurogenic processes were no longer found activated by day 15 p.d. (Fig. 3C; S. Table 4), suggesting that the progression of denervation-activated muscle glial cells toward nerve repair was interrupted from day 5 to day 15. We further investigated the neurogenic progression of muscle glial-derived cells following denervation, by pseudo-time analysis, using Monocle3 (Cao et al., 2019) which infers temporal dynamics of gene expression as cells progress through timepoints. This analysis revealed that muscle glial-derived progenies diverged into two pseudo-temporal trajectories during our experimental conditions, at days 2 and 5 p.d.; however, by day 15 p.d. most of the muscle glial-derived cells collapsed into one common central trajectory (Fig. 3D-F). Module analysis of the DE genes within the pseudo-temporal trajectories could define 8 distinct functional modules (Fig. S5A). According to IPA analysis, module 4, which was more represented in the trajectory of cells at day 2 p.d., was enriched in genes implicated in metabolic reprogramming toward oxidative phosphorylation - a process associated to general commitment to differentiation. Modules 2 and 3, which were enriched in neurogenic processes, such as axonal guidance, or pathways implicated in neurogenesis, including ILK signalling and VDR/RXR activation, were mostly represented in the trajectory of cells at day 5 p.d. (Fig. S5A-C). However, while module 2 was still represented in the trajectory of cells at day 15 p.d., the same trajectory was dominated by the upregulation of genes implicated in extra-cellular matrix (ECM)-related processes, including fibrosis and acute phase response signalling (Fig. S5A-C).

**Figure 3.**
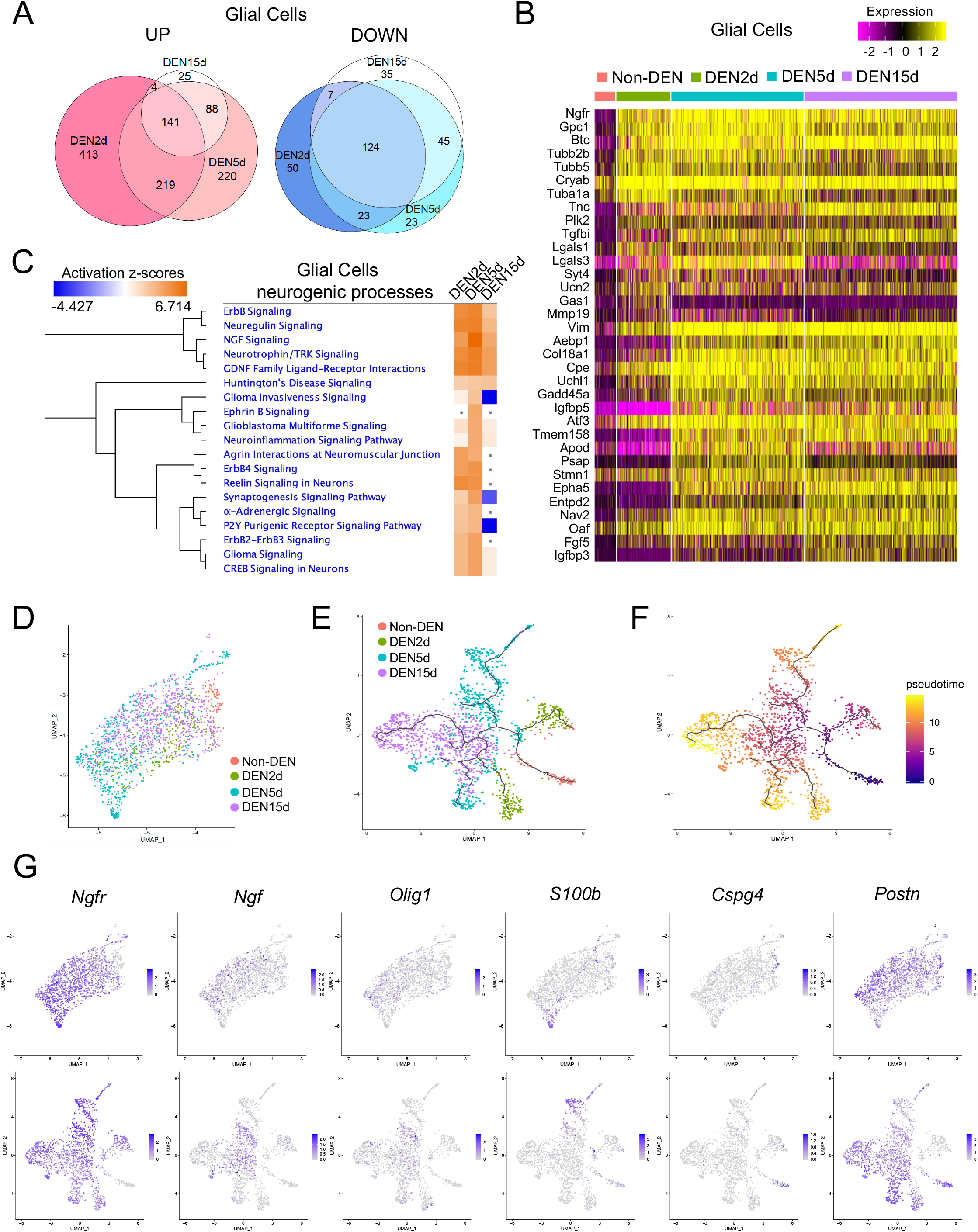
(A) Venn diagram of upregulated genes (left) and downregulated genes (right) in glial cells at DEN2d, DEN5d, and DEN15d compared with Non-DEN. Genes with p-value-adj <= 0.01 are counted. (B) Heatmap of top 20 upregulated genes in glial cells at DEN2d, DEN5d, and DEN15d respectively compared with Non-DEN. (C) IPA comparison of DE genes in glial cells at DEN2d, DEN5d, and DEN15d respectively compared with Non-DEN. Selected pathways related to neurogenic processes are shown. Pathways with p-value >= 0.05 were indicated with gray dots. (D) UMAP embedding subset from the whole dataset of glial cells colored by time point. (E) UMAP embedding of glial cells after reclustering, with superimposition of pseudo-time trajectories from Monocle3. Glial cells are colored by time point. (F) Same as (E), but glial cells are colored by pseudo-time. (G) UMAP embedding of glial cells before (Upper) and after (Bottom) reclustering, showing the expression of selected genes associated with different glial cell-derived lineages.

The comparison of the initial UMAP embedding of the whole dataset with the UMAP embedding of muscle glial-derived cells used for pseudo-time analysis revealed that the cell trajectories identified by Monocle corresponded to distinct glial-derived progenies, whose dynamic gene expression patterns predicted a commitment to specialized cell types in response to denervation (Fig. S5D-E). Indeed, while the glial cell identity marker *Plp1* was invariably expressed by all muscle glial cells in unperturbed muscles as well as at each time point p.d., the majority of neurogenic genes were only induced upon denervation (Fig. S6). Among them, *Ngfr* expression was induced in the large majority of glial-derived cells following denervation (Fig. 3G; Fig. S6). We observed that *Ngfr*-expressing cells aligned toward a gradient of increasing *Ngfr* expression, with higher expression levels observed in cells polarized toward the lower tip of the UMAP that was only detected at day 5 p.d. (Fig. 3G; Fig. S5G). Of note, this lower tip was composed by cells that also expressed presumptive markers of Schwann cells, such as *S100b* (Fig. 3G; Fig. S5D). *S100b* expression also marked a distinct streak of cells in the UMAP (Fig. 3G) that also corresponded to cell trajectories identified by the pseudo-time analysis at days 2 and 5 p.d. (Fig. 3D-F; Fig. S5D-E). Interestingly, by day 5 p.d. a subset of *S100b* expressing cells were aligned into an outstanding streak of that uniformly exhibited a gene expression profile predictive of G2-M phase progression (Fig. S5F), suggesting that they could represent an expansion of Schwann cell progenitors. A recent work has established the identity of a specialized sub-population of Schwann cells within the NMJ, also referred as to Perisynaptic Schwann cells (PSC), that co-express *S100b* and *Cspg4* (NG2) – a gene otherwise expressed in other S100B negative cell types (Castro et al., 2020). However, our data show that co-expression of *S100b* and *Cspg4* could be only detected in glial cells from unperturbed muscles (Fig. 3G; Fig. S6). Thus, the streak of proliferating cells that express *S100b*, but not *Cspg4*, might represent a gradient of Schwann cell progenitors before their functional specialization into PSCs, similar to what observed during development (Castro et al., 2020). The finding that *S100b/Cspg4* expressing cells accounted for the large majority of muscle-resident glial cells in unperturbed muscles but were not detected at any time point p.d. (Fig. S6), suggests an abortive attempt to repair NMJ in denervated muscles, possibly because of the irreversible nature of denervation by complete sciatic nerve section. Moreover, while the large majority of glial cells from unperturbed muscles were expressing *S100b*, in denervated muscles *S100b* was expressed by a fraction of the expanding population of glial-derived cells that was only detected by day 5 p.d. This hints at the possibility that glial cells of unperturbed muscles are mostly composed by Schwann cells, whereas denervation-activated muscle glial cell progenies are more heterogeneous and might contain different populations of cell types in dynamic transition along the time points p.d. (Fig. S5G). For instance, within the high *Ngfr*-expressing cells we identified a subset of cells that also expressed *Ngf* in response to denervation, in concomitance with the appearance of *Olig1* expression (Fig. 3G). These events have been previously associated to Schwann cell de-differentiation, as an early response to nerve injury that typically precedes their expansion and differentiation into Schwann cells toward nerve repair (Jessen and Arthur-Farraj, 2019). Consistently, these cells were polarized toward a cluster of high *Ngfr*-expressing cells at day 5 p.d.; however, by day 15 p.d. these cells re-dispersed among the other muscle glial-derived cells (Fig. 3G; Fig. S5 and S6). A third distinct group of denervation-activated glial cells was marked by the expression of *Postn* (Fig. 3G), without co-expression of *Ngfr*, and was enriched in genes implicated in ECM remodeling, including *Postn* itself, *Tnc, Vim and Itgb8* (Fig. S5D-E and S6) as well as *Serpine1*, *Col1a1* and *Timp1*. This subset coincided with the cell trajectories identified by pseudo-time analysis at days 2 and 5 p.d. that were found in the opposite direction of the cell trajectories enriched with *Ngfr* at the same time points (Fig. 3D-F; Fig. S5D-E). This population progressively expanded along with the time points p.d. and represented the majority of muscle glial-derived cells at day 15 p.d. Interestingly, while these three putative sub-populations of muscle glial-derived cells could be clearly separated, as distinct clusters/trajectories of cells during days 2 and 5 p.d., by day 15 p.d. most of the muscle glial-derived cells repositioned into a diffuse and heterogeneous population of *Postn*-expressing cells (Fig. 3G; Fig. S5D-G; Fig. S6). Overall, the trend of DE genes in muscle glial-derived cells along with our timepoints post denervation suggests an initial activation of pro-neurogenic processes toward nerve regeneration and recovery of NMJs that eventually fails and is followed by the retention of a heterogeneous population of glial-derived cells expressing genes implicated in ECM remodelling.

#### Activated Fibroblasts

Activated fibroblasts represent a population of mesenchymal cells that share many gene expression features with FAPs, the most contiguous cell type, as shown by the UMAP embedding (Fig. 1B). While both activated fibroblasts and FAPs express *Cd34*, a cell surface glycoprotein that is generally expressed in hematopoietic and mesenchymal muscle-resident cells (Sidney et al., 2014), lower expression of *Pdgfra* and *Ly6a* could discriminate activated fibroblasts as a cellular cluster independent from FAPs (Fig. S2). Moreover, at variance with FAPs, which represent the most abundant fraction of muscle-resident cells, a modest amount of activated fibroblasts (2.2% of the whole dataset) was detected in unperturbed muscles (Fig. 2A and S. Table 1); however, similar to muscle glial cells, activated fibroblasts progressively expanded following denervation (S. Table 1; Fig. S3A; see also Fig. 4G) and exhibited significant changes in gene expression between days 2 and 5 p.d., with a strong bias toward gene upregulation (S. Table 2; Fig. 4A and B). IPA analysis of DE genes at each timepoint p.d. predicted the consensual activation of several signaling pathways implicated in neurogenic processes related to synaptogenesis, axon guidance and neuronal migration, among the others (Fig. 4C).

**Figure 4.**
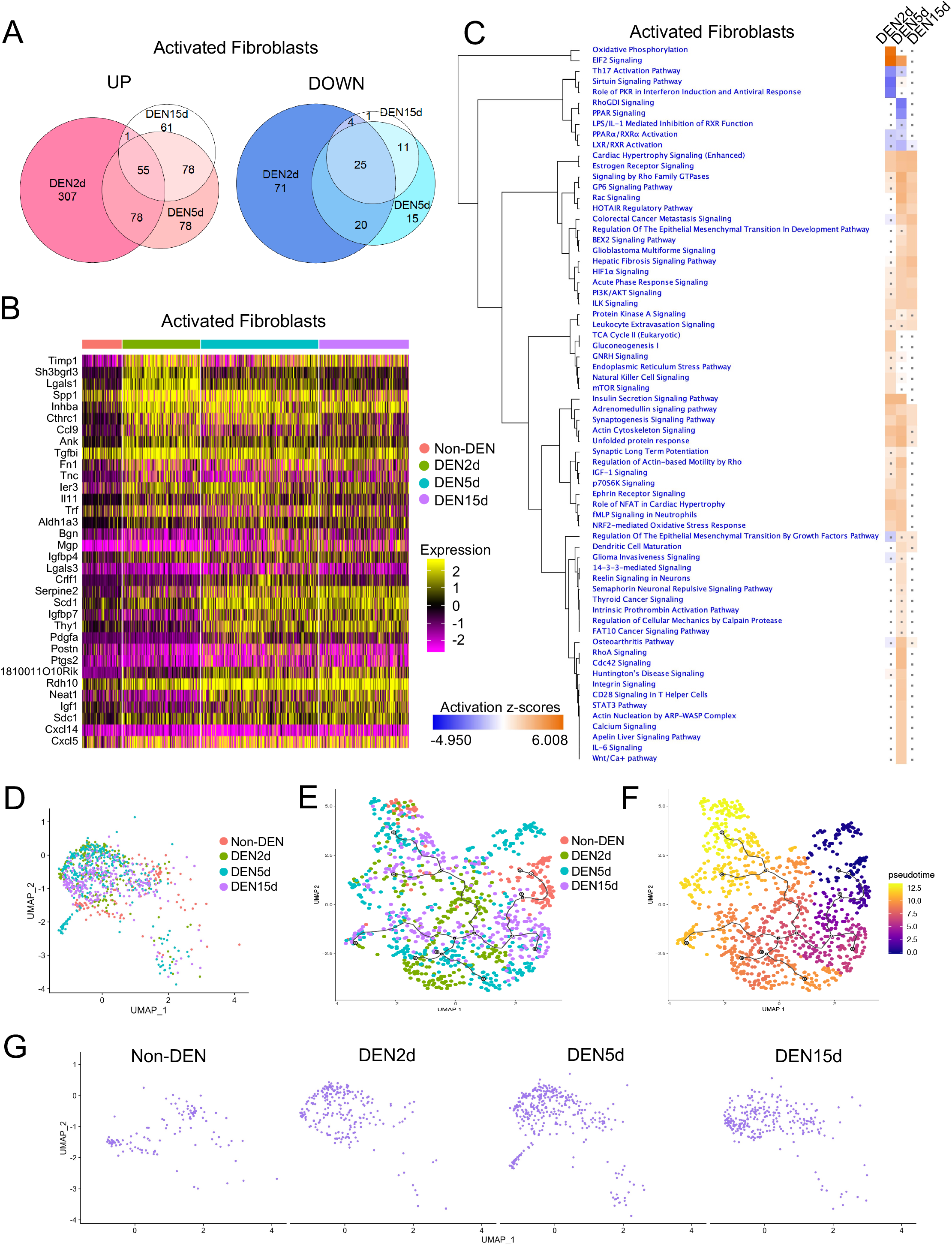
(A) Venn diagram of upregulated genes (left) and downregulated genes (right) in activated fibroblasts at DEN2d, DEN5d, and DEN15d compared with Non-DEN. Genes with p-value-adj <= 0.01 are counted. (B) Heatmap of top 20 upregulated genes in activated fibroblasts at DEN2d, DEN5d, and DEN15d respectively compared with Non-DEN. (C) IPA comparison of DE genes in activated fibroblasts at DEN2d, DEN5d, and DEN15d respectively compared with Non-DEN. Pathways significantly enriched in either time point with absolute z score > 1 were shown. Pathways with p-value >= 0.05 were indicated with gray dots. (D) UMAP embedding subset from the whole dataset of activated fibroblasts colored by time point. (E) UMAP embedding of activated fibroblasts after reclustering, with superimposition of pseudo-time trajectories from Monocle3. Activated fibroblasts are colored by time point. (F) Same as (E), but here activated fibroblasts are colored by pseudo-time. (G) UMAP embedding of activated fibroblasts before reclustering in time-lapse across denervation time points.

Interestingly, similar to muscle glial cells, also in activated fibroblasts these neurogenic signals were downregulated by day 15 p.d., while DE genes implicated in ECM remodeling toward cell proliferation, migration and fibrosis (such as *Serpine 2, Scd1, Igfbp7* and *Thy1*) continued to be expressed (Fig. 4B). While pseudo-time analysis could not identify clear cellular trajectories in activated fibroblasts along with the time points p.d. (Fig. 4D-F), comparative analysis with UMAP indicated a general trend of activated fibroblasts to form sub-clusters at days 2 and 5 p.d., followed by a progressive dispersion within the heterogenous main cluster observed by day 15 p.d. (Fig. 4D-G).

Overall, the consensual activation pattern of muscle glial cells and activated fibroblasts suggests that these two denervation-activated muscle resident cell types might reciprocally exchange neurogenic signals toward repair of NMJs and possibly nerve regeneration. The interruption of this neurogenic signaling network by day 15 p.d. in both cell types suggests a functional interdependence between muscle glial cells and activated fibroblasts in promoting nerve repair. At the same time, their simultaneous conversion into cellular sources of pro-fibrotic ECM components by day 15 p.d. also suggests that these cell types might cooperate to promote persistent fibrosis in chronically denervated muscles.

#### Other muscle-resident cell types and inflammatory cells

We have detected modest changes in number and DE genes in other muscle-resident cell types.

FAPs represent the largest population of muscle-resident cells and have been previously identified as source of pro-atrophic and pro-fibrotic signals in denervated muscles (Madaro et al., 2018). Our data show that although FAPs exhibited modest changes in gene expression that mostly occurred by day 5 p.d. (Fig. 5A and B), a reconfiguration of sub-clusters (Fig. S3A; Fig. S7) coincided with the progressive activation of pro-fibrotic gene networks (Fig. 5A-C). Interestingly, genes implicated in synaptogenesis and other presumptive glial/neurogenic signaling (BEX2 and ILK signaling) were also activated in FAPs at late time points p.d. (Fig. 5A-C). SNN clustering identified four discrete FAPs sub-clusters (Fig. S7A), which were marked by relative enrichment of genes previously assigned to specific sub-populations of FAPs (Malecova et al., 2018; Oprescu et al., 2020). Three of these sub-clusters (namely, number 1, 3 and 4) were identified by the relative enrichment in the expression of genes previously associated with FAPs quiescence and propensity to adopt an adipogenic phenotype (i.e., *Dpp4* and *Tek* in sub-cluster 3) (Fig. S7A and D) (Malecova et al., 2018; Merrick et al., 2019). These three sub-clusters accounted for most of the FAP population present in unperturbed muscles (Fig. S7B-C). Upon denervation, sub-cluster 2, which was enriched in genes associated with FAP activation following injury, such as *Vcam1, Serpine1, Timp1* and *Lif* (Malecova et al., 2018; Oprescu et al., 2020) progressively expanded and persisted until day 15 p.d. (Fig. S7B-D). Of note, sub-cluster 2 also contained a subset of denervation-activated FAPs exhibiting progressive activation of STAT3-IL6 signaling (Fig. S7C-D), which promotes myofiber atrophy and muscle fibrosis (Madaro et al., 2018). High levels of *Il6* were also detected by other muscle-resident cell types in close proximity with FAPs, such as activated fibroblasts and pericytes during the time points p.d. (Fig. S2). When compared to the kinetic of FAPs activation by myotrauma, FAPs from denervated muscles differed in term of magnitude of activation (much less expansion and DE genes), but also in term of recovery post-perturbation, as even by day 15 p.d. the original configuration of FAP sub-clusters typical of unperturbed muscles was not resumed (Fig. S7B-C).

**Figure 5.**
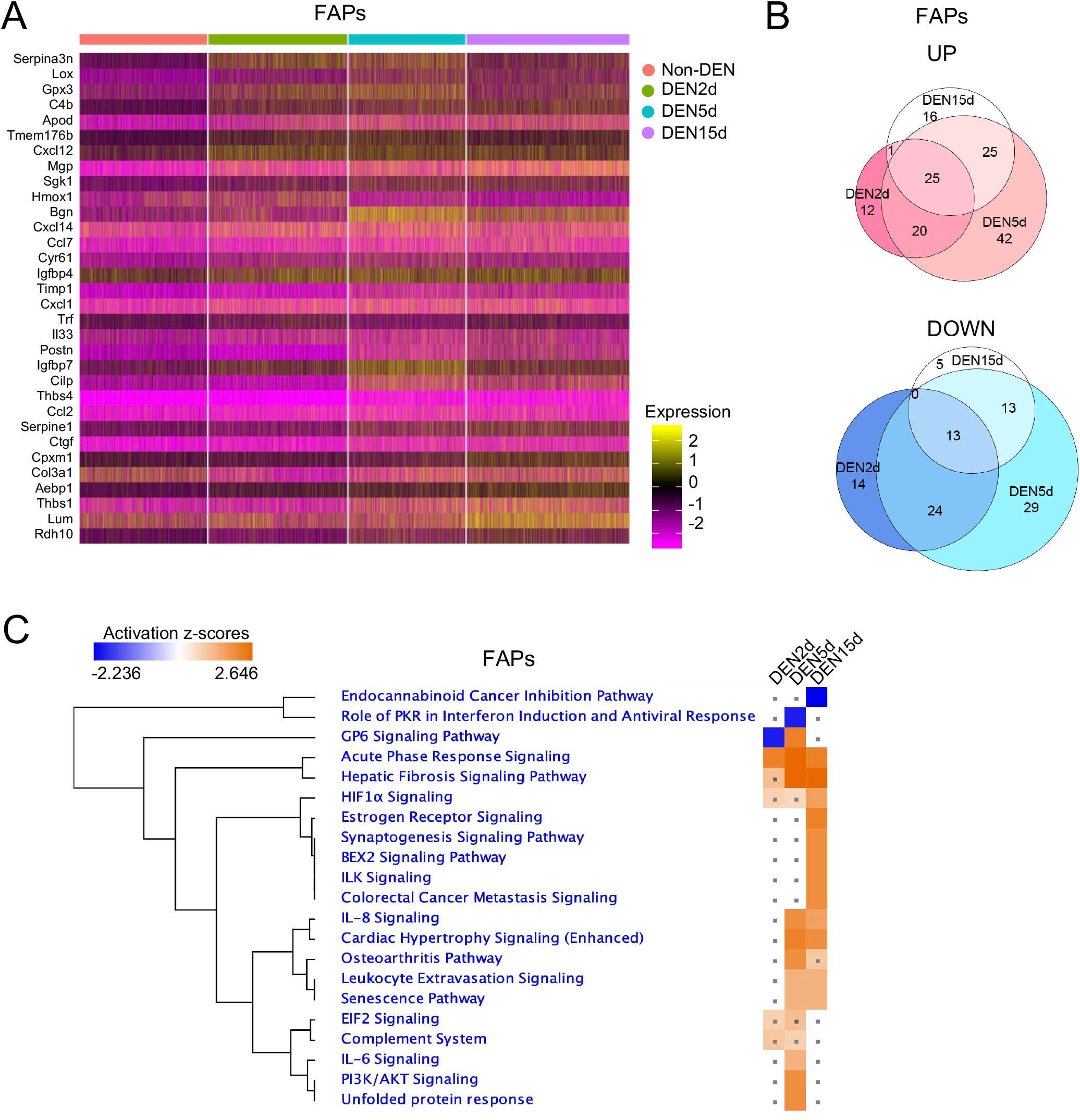
(A) Heatmap of top 20 upregulated genes in FAPs at DEN2d, DEN5d, and DEN15d respectively compared with Non-DEN. (B) Venn diagram of upregulated genes (Upper) and downregulated genes (Bottom) in FAPs at DEN2d, DEN5d, and DEN15d compared with Non-DEN. Genes with p-value-adj <= 0.01 are counted. (C) IPA comparison of DE genes in FAPs at DEN2d, DEN5d, and DEN15d respectively compared with Non-DEN. Pathways significantly enriched in either time point with absolute z score > 1 were shown. Pathways with p-value >= 0.05 were indicated with gray dots.

Vessel-associated muscle-resident cells include endothelial cells, SMMCs and pericytes. SMMCs and pericytes exhibited negligible fluctuation in number and gene expression during the time points p.d. (S. Table 1 and 2), while endothelial cells showed a transient drop in cell number by day 2 p.d., followed by a recovery between day 5 p.d. (S. Table 1). Accordingly, most of DE genes were detected between day 2 and 5 p.d. (S. Table 2) and predicted an activation of signaling implicated in different processes, such as cell growth, senescence, oxidative stress response, neo-angiogenesis and fibrosis (Fig. 6A-C).

**Figure 6.**
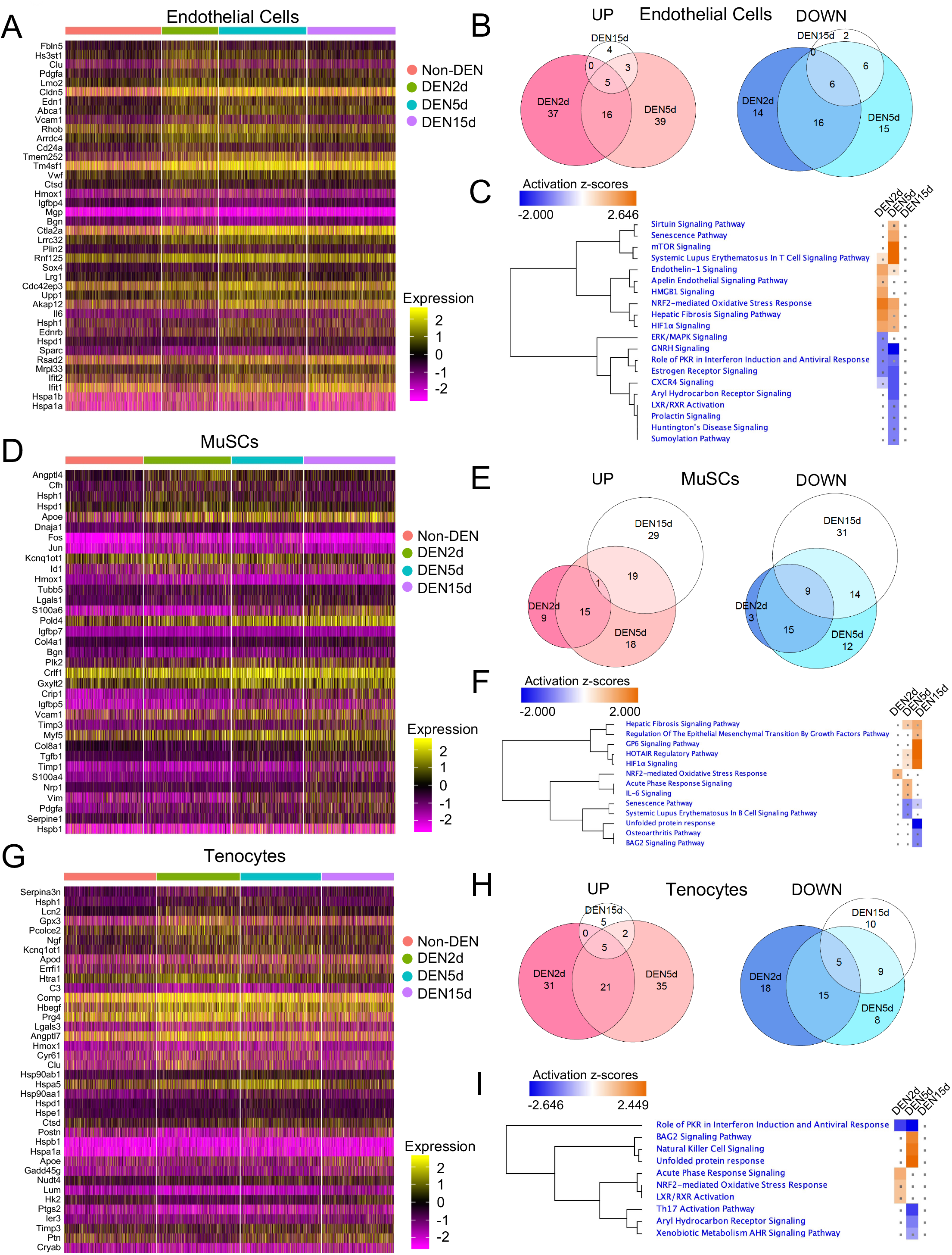
(A, D, G) Heatmap of top 20 upregulated genes in endothelial cells (A), MuSCs (D), and tenocytes (G) at DEN2d, DEN5d, and DEN15d respectively compared with Non-DEN. (B, E, H) Venn diagram of upregulated genes (left) and downregulated genes (right) in endothelial cells (B), MuSCs (E), and tenocytes (H) at DEN2d, DEN5d, and DEN15d compared with Non-DEN. Genes with p-value-adj <= 0.01 are counted. (C, F, I) IPA comparison of DE genes (P-value-adjusted < 0.01) in endothelial cells (C), MuSCs (F), and tenocytes (I) at DEN2d, DEN5d, and DEN15d respectively compared with Non-DEN. Pathways significantly enriched in either time point with absolute z score > 1 were shown. Pathways with p-value >= 0.05 were indicated with gray dots.

MuSCs and tenocytes did not show significant changes in cell number during the time points p.d. (S. Table 1) and exhibited modest changes in gene expression (S. Table 2; Fig. 6D-I). The few genes induced by denervation in MuSCs converged on a general trend of activation of fibrosis and epithelial-mesenchymal transition (EMT) and response to hypoxia (Fig. 6F). The lack of evidence of MuSC activation by denervation is in apparent conflict with the activation of MuSCs following sciatic nerve transection reported by Liu and colleagues (Liu et al., 2015b); however, we noticed that by day 2 p.d. a small fraction of MuSCs transiently expressed *Stmn1,* which was also expressed in muscle glial-derived cells (Fig. S8). Interestingly, by day 5 p.d. *Stmn1*-expressing cells aligned along a streak of cells in linear contiguity with denervation-activated *S100b* expressing muscle glial-derived cells (Fig. S8). This fraction of *Stmn1*-expressing cells connected MuSCs and muscle glial-derived cells and was likely composed of proliferating cells, as suggested by the high expression of the proliferation marker *Mki67* and cyclins that induce cell cycle progression, such as *Ccna2*, *Ccnb1*, *Ccnb2* and *Ccnd1* (Fig. S8). Moreover, we detected a number of common DE genes in the streak of cells connecting MuSCs and muscle glial-derived cells (S. Table 5), suggesting that they could be a population of cells transitioning between these two lineages. Co-expression of *Stmn1* and *Mki67* has been recently used to identify adult gastric stem cells (Han et al., 2019) and STMN1 deficiency has been associated to axonopathy of peripheral nerves in age-dependent manner (Liedtke et al., 2002). Future work will be required to determine the identity and function of these cells.

Finally, denervation did not trigger an immediate and massive infiltration by inflammatory cells, with related changes in gene expression, that were typically observed in response to myotrauma (De Micheli et al., 2020; Dell’Orso et al., 2019; Oprescu et al., 2020; Pawlikowski et al., 2019; Petrany et al., 2020). Most of DE genes were detected by day 2 p.d., with more downregulated genes; by contrast, by day 5 p.d. the trend was inverted, with the large majority of DE being upregulated (Fig. 7A and B). While the DE genes annotated at day 2 p.d. were implicated in a variety of processes related to metabolism, inflammation/immune response and extra-cellular signaling pathways, by day 5 p.d. only gene implicated in the activation of signaling promoting dendritic cell/macrophage maturation and pro-fibrotic signals could be detected (S. Table 6). SNN clustering identified 3 major sub-clusters (Fig. 7C and D), that were mostly unaltered during the timepoints p.d., although persistence of macrophages accumulation was observed by day 15p.d. (Fig. 7E). This general trend of gene expression indicates a mild activation of an immune response at early stages post-denervation, but also suggests a lack of complete resolution of the inflammation, leading to the persistence of inflammatory cells that can potentially contribute to fibrosis.

**Figure 7.**
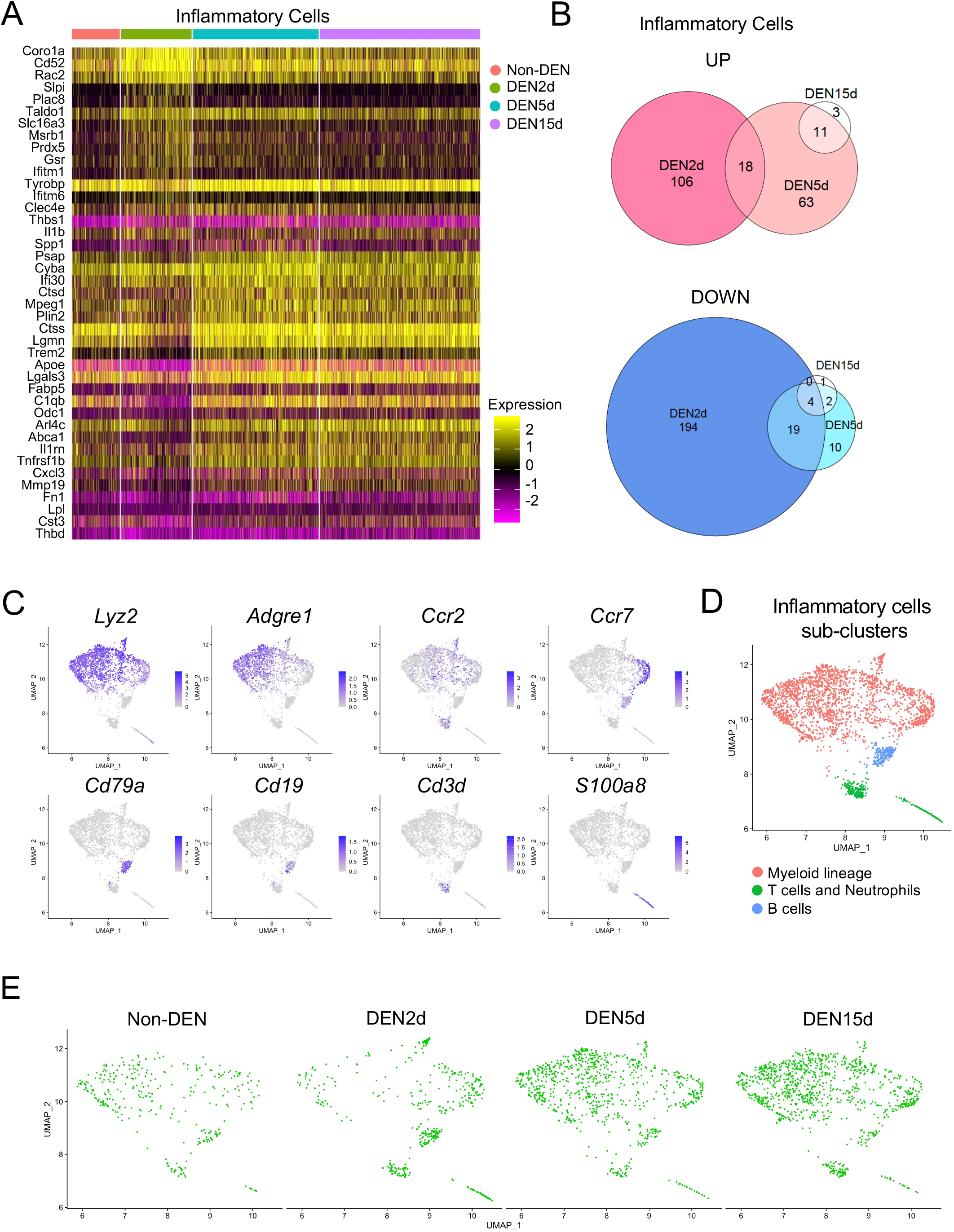
(A) Heatmap of top 20 upregulated genes in inflammatory cells at DEN2d, DEN5d, and DEN15d respectively compared with Non-DEN. (B) Venn diagram of upregulated genes (Upper) and downregulated genes (Bottom) in inflammatory cells at DEN2d, DEN5d, and DEN15d compared with Non-DEN. Genes with p-value-adj <= 0.01 are counted. (C) UMPA embedding of inflammatory cells showing the expression of selected genes associated with sub-clusters including macrophages, T cells, neutrophils, and myeloblasts. (D) UMAP of sub-clusters within inflammatory cells. (E) UMAP embedding of inflammatory cells in time-lapse across denervation time points.

## Discussion

Our longitudinal analysis of the transcriptome from denervated skeletal muscles at the single cell level provides an unprecedented atlas of the transcriptional response of skeletal muscle-resident cells to acute denervation. Our analysis identified muscle-glial cells and activated fibroblasts as the main muscle-resident cell types activated in response to denervation. Their gene expression profiles predict that they are also the cellular sources of muscle-derived signals elicited by complete transection of the sciatic nerve.

Our scRNA-seq data show that, at variance with the dramatic expansion of multiple cell types observed upon acute muscle injury by myotrauma, the transcriptional perturbations in muscle-resident cell types following acute denervation are much less extensive, while specifically directed toward the selective activation of two cells types implicated in nerve repair. Indeed, denervation-activated muscle-glial cells and activated fibroblasts exhibited discrete expression patterns of genes implicated in the regeneration of cellular and extracellular components of NMJ (e.g. Schwann cells and ECM components of NMJ), as well as neurotrophic and axon guidance signals, consistent with their functional interplay during peripheral nerve regeneration (Parrinello et al., 2010).

Our data show that while *S100b*-expressing Schwann cells account for a large amount of muscle-resident glial cells in unperturbed muscles, after denervation only a small proportion of glial-cell derived progenies (less than 10%) expressed *S100b* by day 15 p.d. This is an indirect evidence of de-differentiation of Schwann cells post-denervation to give rise to multiple neurogenic lineages, as described in other models of neurogenic regeneration post nerve injury (Gordon, 2020). However, this initial pro-neurogenic response ceased at later stages post-denervation, indicating an abortive attempt to regenerate denervated muscles.

Although larger populations of muscle-resident cell types (e.g. FAPs, endothelial and inflammatory cells) exhibited modest changes in gene expression and size in response to denervation, as compared to muscle injury, we observed reconfigurations of sub-clusters within these cell types along the all timepoints p.d. Sub-clusters within heterogenic muscle-resident cell types, such as FAPs, likely reflect cellular states in dynamic transition in response to muscle injury (Malecova et al., 2018). The observation that denervation-induced sub-clusters of FAPs and endothelial cells enriched in genes implicated in ECM remodeling and fibrosis persisted through all the timepoints p.d. suggests a general lack of resolution of denervation-activated processes that possibly contributes to the progressive fibrosis typically observed in denervated muscles. In particular, it is possible that denervation-activated mesenchymal cells initially produce specific ECM components of the NMJ-associated basal lamina (Singhal and Martin, 2011), but upon lack of completion of NMJ repair these cells might continue to constitutively produce ECM components ultimately leading to fibrosis. Thus, despite the different spectrum of cell types implicated, skeletal muscle responses to denervation or myotrauma share interesting analogies. For instance, in both cases the cell types activated are finalized to specific biological processes – i.e. MuSC-mediated regeneration of myofibers or muscle glial-derived generation of NMJ cellular components (e.g. Perisynaptic Schwann cells). And in both cases, other mesenchymal cell types, such as activated fibroblasts or other interstitial cells, provide regulatory signals and changes in ECM that favor myofiber or NMJ regeneration. However, these signals are typically terminated upon successful regeneration, possibly by feedback signals derived from repaired structures. In our experimental model of complete sciatic nerve transection, the denervation is irreversible and the failure to repair nerve and NMJ likely causes the lack of resolution of denervation-activated processes that ultimately culminates with chronic fibrosis. Conceivably, upon complete sciatic nerve transection, lack of signals from motor neurons and loss of coordination between denervation-activated cell types might lead to a persistent pathological reconfiguration of ECM composition. This outcome resembles the fibrotic degeneration associated to chronic degenerative muscular disorders that impairs muscle regeneration (Farup et al., 2015).

Overall, our data predict a functional interdependence between denervation-responsive cell types and identify networks implicated in both pathogenic events (myofiber atrophy and fibrosis) and nerve regeneration that could be exploited for interventions toward promoting nerve repair and limiting muscle degeneration and atrophy. These data will provide an initial atlas of reference for further studies in experimental conditions of reversible nerve damage, in order to capture cellular networks that can be targeted by interventions toward promoting nerve repair. Our data could also guide the selection of genes that discriminate specific cell types or sub-clusters activated by denervation for future strategies of FACS-mediated sorting or genetic model of lineage tracing of specific cell types of interested.

## Acknowledgments

We thank Brian James for library preparation and Amy Cortez for assistance on FACS-mediated isolation of single cells. We are also grateful to Drs Vittorio Sartorelli, Lorenzo Giordani, Mattia Forcato and Alessandra Sacco for critical reading of the manuscript and helpful discussion during the execution and analysis of the experiments.

This work was supported by NIH/NIAMS grant (R01AR076247-01) to PLP; American Heart Association Postdoctoral Fellowship (19POST34450187) to CN; Italian Ministry of Health (GR-2013-02356592) and “Roche per la Ricerca 2019” grant to L.M.

## Author Contributions

CN performed scRNA-seq data analysis and contributed to the experimental design, manuscript writing and figure preparation. XW contributed to data analysis and interpretation, manuscript writing and figure preparation; UE contributed to the experimental design and scRNA-seq sample preparation; DP and LM contributed to the data interpretation by sharing datasets and contributed manuscript writing and figure preparation. PLP designed the experiments, contributed to the data analysis and interpretation, and wrote the manuscript.

## Methods

### Animals

All experiments in this study were performed in accordance with protocols approved by the Sanford Burnham Prebys Medical Discovery Institute (SBP) Animal Care and Use Committee (IACUC) and the Italian Ministry of Health, National Institute of Health (IIS) and Santa Lucia Foundation (Rome). The study is compliant with all relevant ethical regulations regarding animal research. C57BL/6J mice were provided by the SBP Animal Facility (La Jolla, CA, USA).

### Muscle denervation

Muscle denervation was achieved by transecting 5 mm sciatic nerve near to the head of the femur of male C5BL6/N mice under anaesthesia. Detailed procedures were described in previous work (Madaro et al., 2018).

### Sample processing for single-cell RNA-sequencing

For each time-point post denervation (namely, 2, 5 and 15 days p.d., abbreviated as DEN2d, DEN5d and DEN15d in all figures and tables) as well as the non-denervated control (abbreviated as Non-DEN in all figures and tables), 1 wild-type male C5Bl6/N mice at 3 months of age was used for each of the two biological replicates. After sciatic nerve transection, we collected two muscles downstream to the sciatic nerve, the tibialis anterior (TA) and gastrocnemius (GA) muscles. Samples were processed as previously described in (Liu et al., 2015), and sorted by fluorescence activated cell sorting (FACS) to remove cell debris, doublets and dead cells. 100,000 live cells/sample were sorted and used for downstream processing with the 10X Genomics 3’ v3 kit for single-cell gene expression. 12,000 cells were used for library preparation, a little over the 10,000 suggested by the 10X protocol, in order to reach a higher capture rate. Libraries were constructed per manufacturer’s instructions and sequenced using Illumina’s NextSeq platform. Average number of cells across samples was 6,000 and average read depth was 44,500 reads/cell. Reads were then aligned to the mouse genome mm10/GRCm38 using CellRanger (v3.1.0). Doublets presence for each replicate was inferred with Scrublet (v0.2.1) (Wolock et al., 2019) within Python (v3.8). Filtered barcode and count matrices produced by CellRanger were used for downstream analyses in R (v3.6.2).

### Data preprocessing, dimensionality reduction and visualization

Seurat (v3.1.2) was used for data filtering, normalization, differential marker expression and visualization (Stuart et al., 2019). Biological replicates for each time point were initially merged together and filtered for quality control parameters (cells with more than 20% reads mapping to mitochondrial genes, less than 200 or greater than 6,000 features (i.e., genes), and more than 50,000 reads were filtered out, as well as cells with a doublet score, derived from Scrublet, greater than 0.1). Data from different time points were then merged, and Seurat’s SCTransform function was used to normalize and scale the data to minimize batch effects (Hafemeister and Satija, 2019), normalizing counts for mitochondrial genes content, number of features and cell cycle phase. Dimensionality reduction was performed through Principal Component Analysis (PCA) on top variable features, with npcs=100. FindNeighbors, FindClusters and UMAP embedding functions were based on the top 20 PCs of the PCA. Clusters were identified through FindClusters with parameters resolution= 0.4, algorithm= 3. UMAP embedding was run with metric= ‘euclidean’.

### Differential marker expression and cell type identification

Identified cell clusters were evaluated for marker gene expression to determine cell types through Seurat’s FindAllMarkers function. Meta-clusters were manually annotated based on shared enriched markers expression of genes reported in literature as cell type specific markers. The FindMarkers function was used to derive differential marker expression per time point, where each time point post-denervation was compared to the non-denervated condition.

### Cell type-specific analysis

We used Seurat’s subset function to extract cells belonging to each meta-cluster, on which we performed new clustering and dimensionality reduction, keeping the same parameters used for the analysis of the complete dataset. The subset data was used for trajectory inference with Monocle.

### Trajectory inference and enriched gene modules analysis

Monocle (v3.0.2.1) (Cao et al., 2019) was used for pseudotime analysis in glial cells and activated fibroblasts and to find enriched gene modules along the pseudotime trajectory of glial cells. Raw data from glial cells and activated fibroblasts as well as their meta-data were used as input for the new_cell_data_set function, to create Monocle objects for trajectory inference. The preprocess_cds function was used to perform PCA analysis and normalize the data, with num_dim = 100, while UMAP embedding was inherited from the original cell population specific Seurat object. Also, gene loadings from the 20 PCs from the cell population specific Seurat object were imported in the Monocle object. For pseudotime inference, the non-denervated time point was chosen as root node.

To identify genes associated to the different cell fates in the glial cells pseudotime trajectory, we calculated differential genes with the graph_test function (with neighbor_graph=“principal_graph”) and selected genes with q-value< 0.000001. These genes were then used to derive gene modules through the find_gene_modules function.

### Pathway analysis

Differentially expressed genes (p-value-adjusted < 0.01) in each cell group at DEN2d, DEN5d, DEN15d respectively compared with Non-DEN were subjected to Ingenuity Pathway Analysis (IPA) (QIAGEN, version 01-16) for “Core Analysis” and further “Comparison Analysis”. Selected pathways, significantly enriched in either DEN2d, DEN5d, or DEN15d with absolute z score >1, were shown in figures and supplementary tables. Pathways with p-value >= 0.05 were indicated with gray dots.

---

**Supplementary Figure 1.**
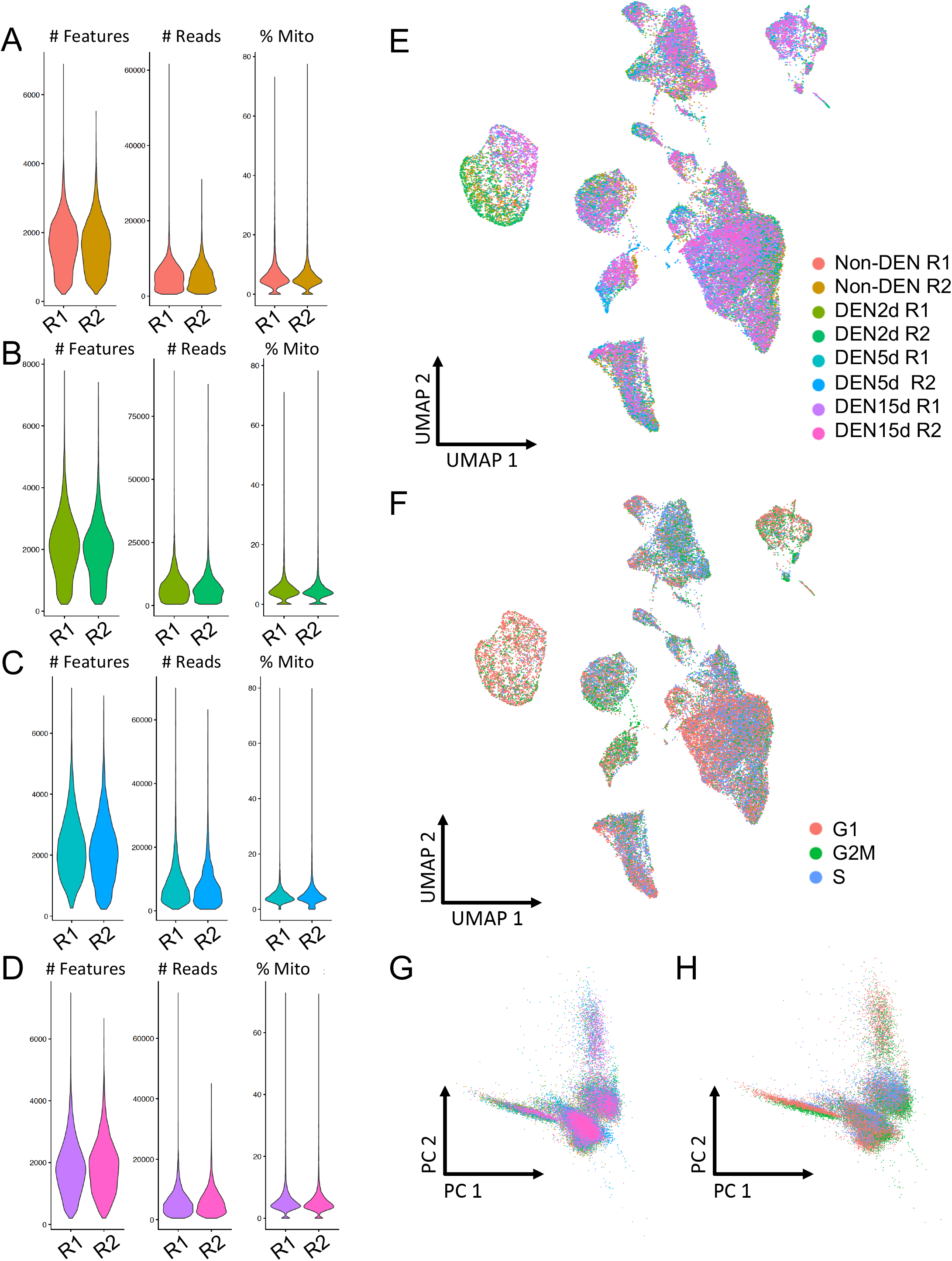
(A-D) Number of genes (# Features), sequencing reads (# Reads) and percentage of mitochondrial transcripts (% Mito) for each cell and each biological sample (R1 and R2) in (A) non-denervated, (B) 2-days, (C) 5-days and (D) 15-days post-denervation. (E) UMAP embedding of all the single cells comprising the dataset colored by biological replicate. (F) UMAP embedding of all the single cells comprising the dataset colored by cell cycle phase. (G-H) PCA embedding of all the single cells comprising the dataset colored by either (G) biological replicate or (H) cell cycle phase.

**Supplementary Figure 2.**
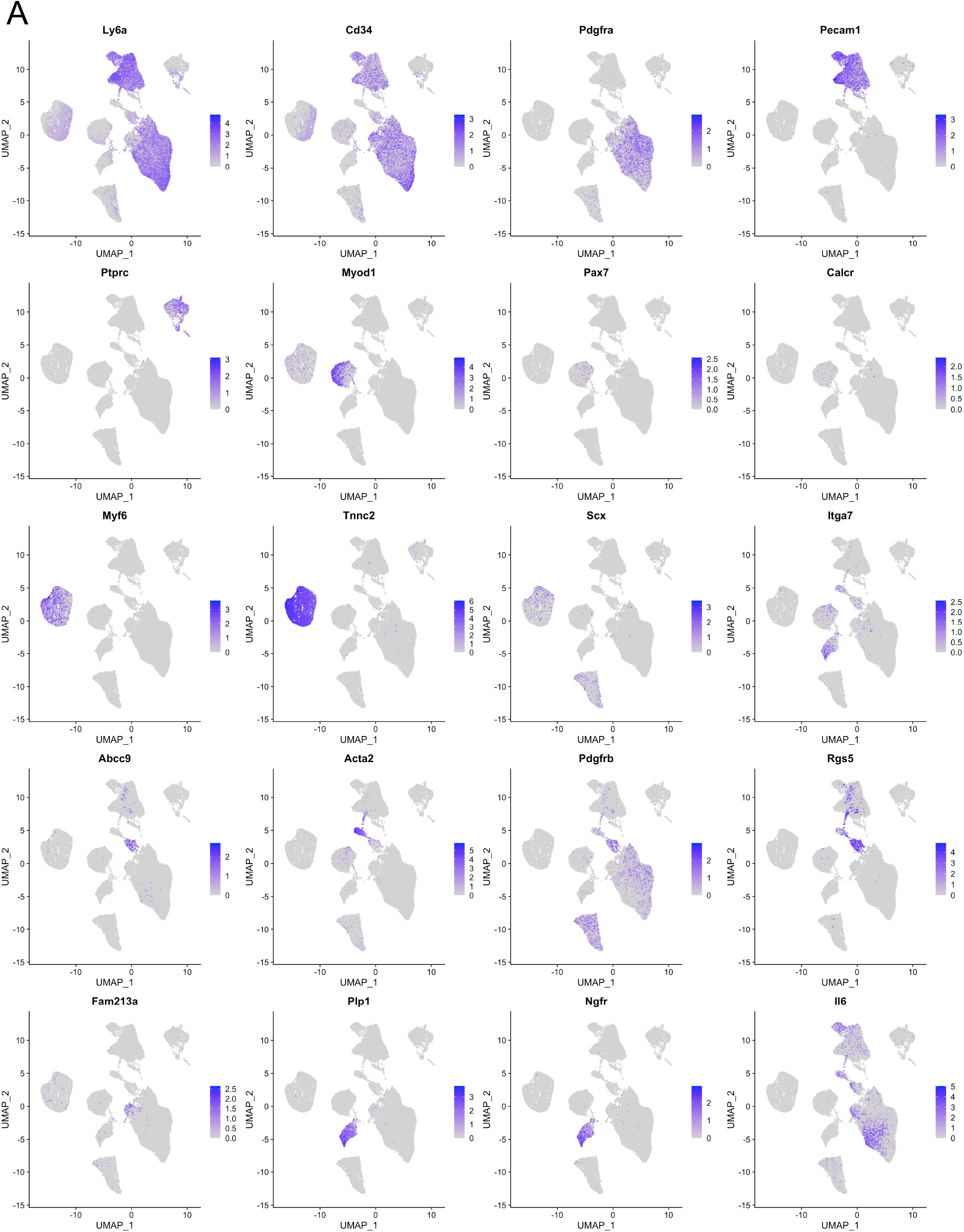
(A) UMAP embedding of all the single cells comprising the dataset showing the normalized expression level of the marker genes in Fig. 1B.

**Supplementary Figure 3.**
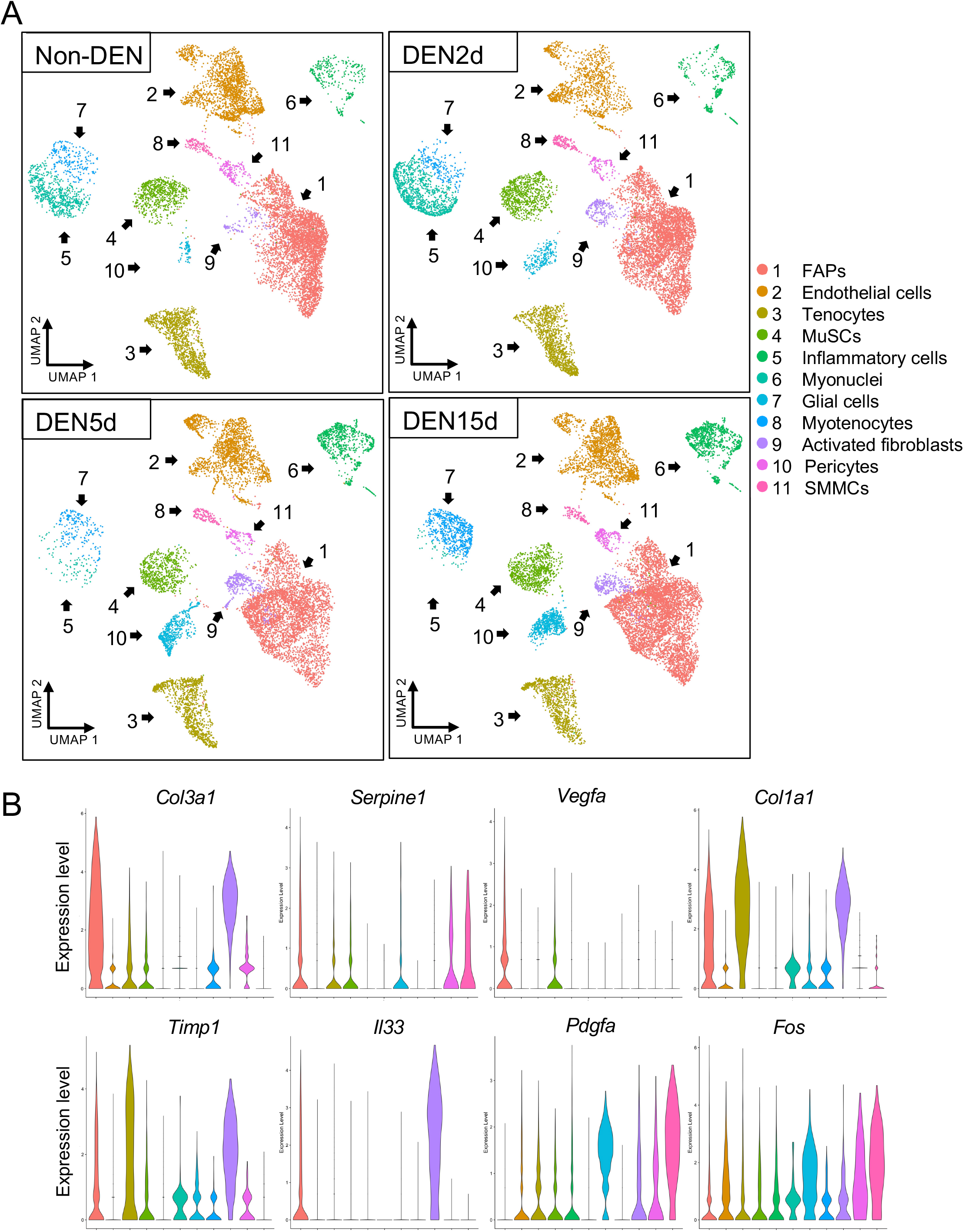
(A) UMAP embedding of single cells at Non-DEN, DEN2d, DEN5d, and DEN15d respectively. (B) Violin plots in DEN5d cells of genes involved in Hepatic Fibrosis Signaling Pathways.

**Supplementary Figure 4.**
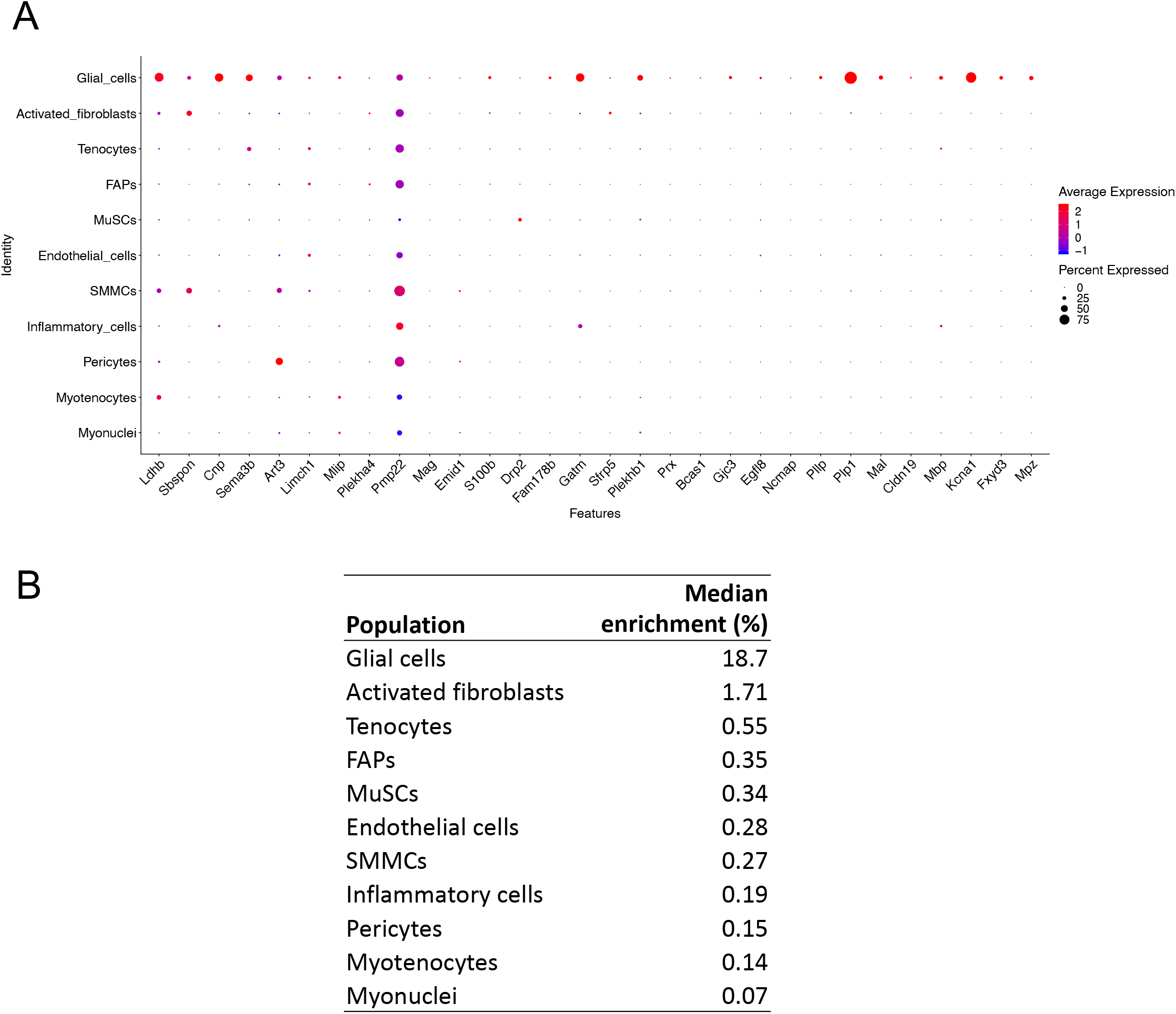
(A) Dot plot of Schwann cells-specific genes extrapolated from earlier scRNA-seq dataset (Pawlikowski et al., 2019) in our cell populations. (B) Table with the median percentage of cell in each population expressing the marker genes in (A).

**Supplementary Figure 5.**
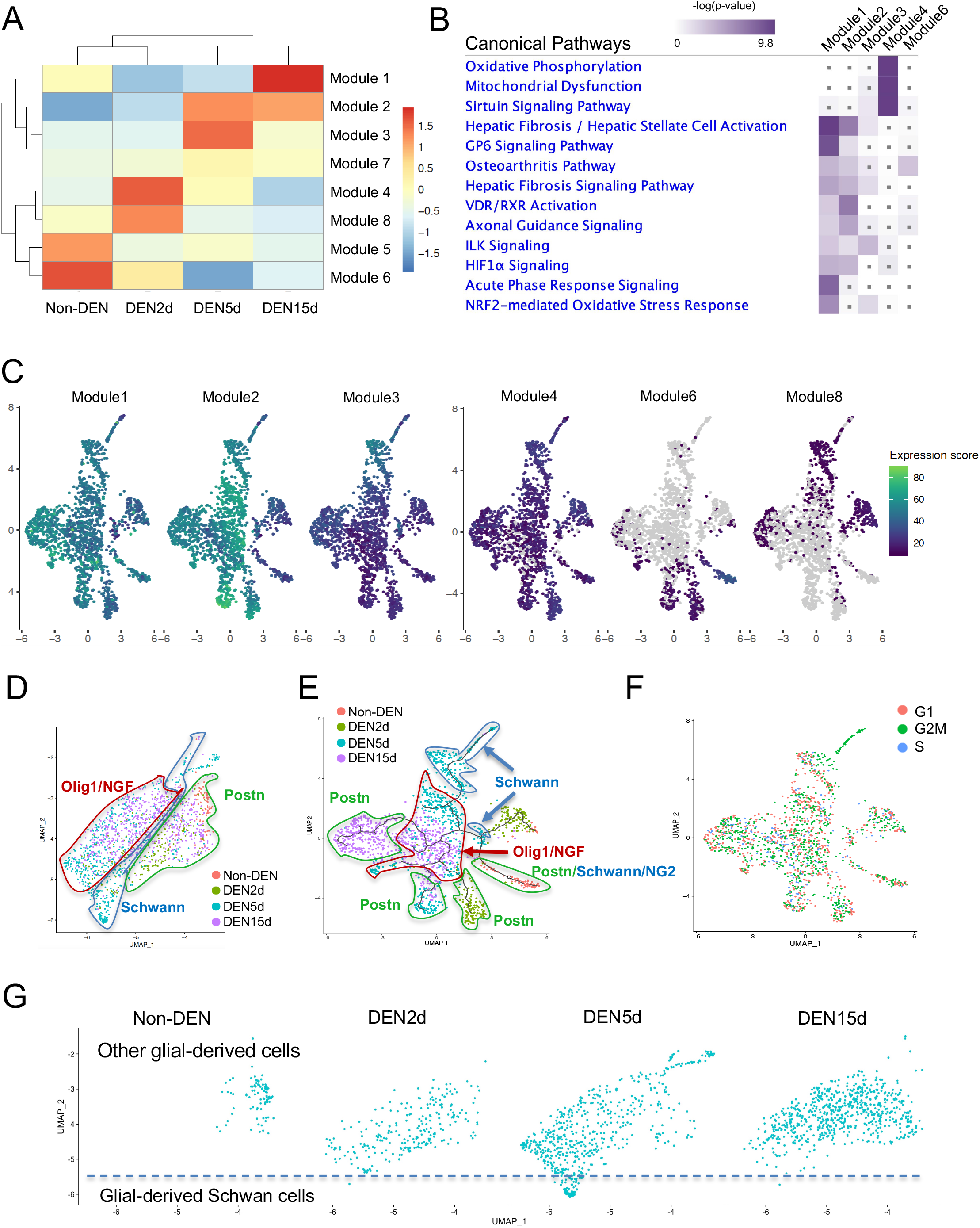
(A) Monocle3 module analysis of co-expression of genes found differentially expressed along the pseudo-time trajectory. (B) IPA analysis of genes belonging to modules expressed at the different experimental time points. (C) UMAP embedding of the enrichment of genes belonging to the major modules. (D, E) Same as Figure 3D and 3E, with circles indicating different glial-derived lineages. Red circle: cells expressing *Olig1* and *Ngf*; Green circle: cells expressing *Postn*; Blue circle: cells expressing Schwann cells genes. (F) UMAP embedding of glial cells after reclustering, colored by cell cycle phase. (G) UMAP embedding of glial cells before clustering in time-lapse across denervation time points. Dash line delineates the lower tip of denervation-activated muscle glial cells that is specific of day 5 p.d.

**Supplementary Figure 6.**
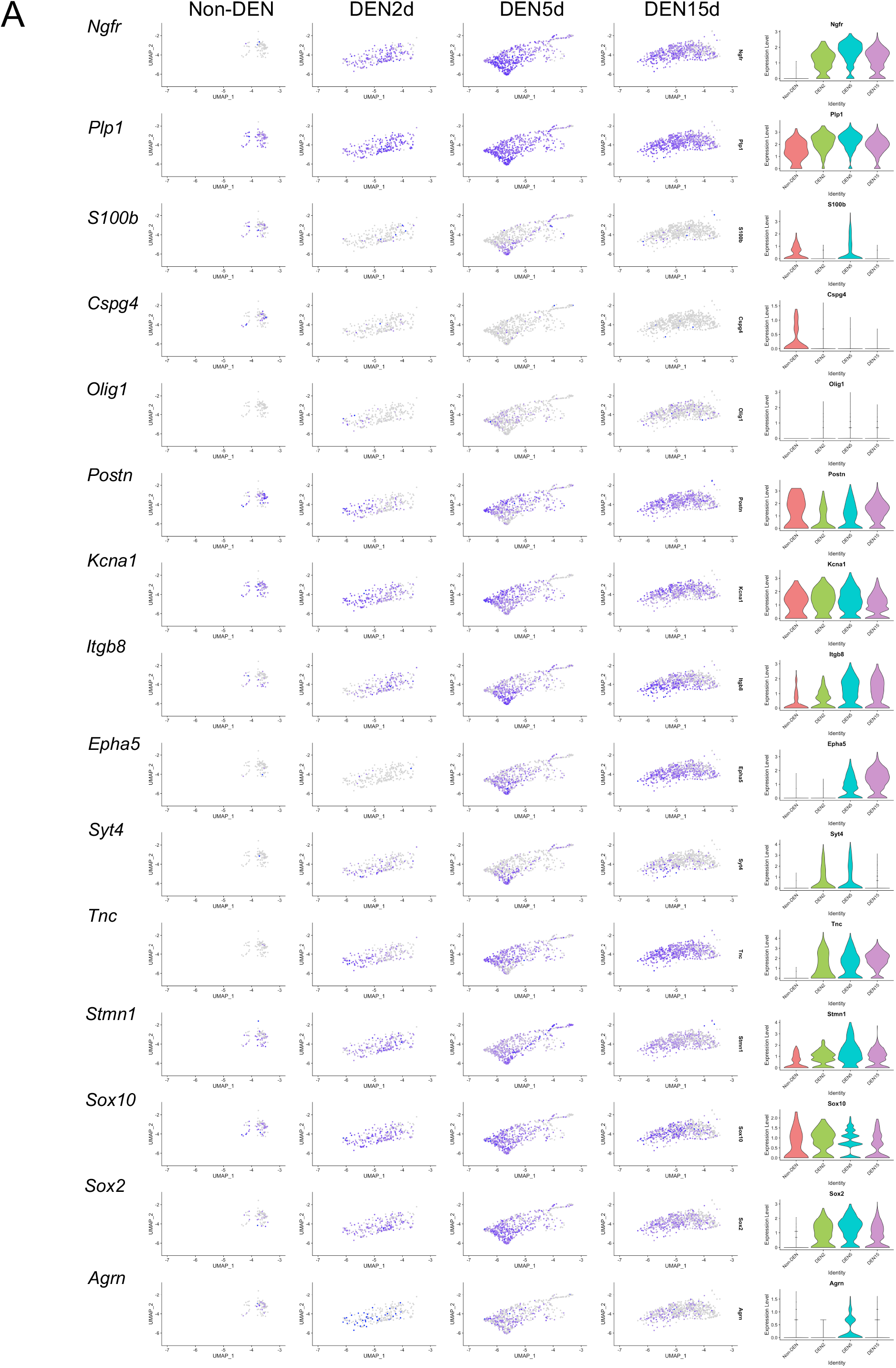
(A) UMAP (left) and Violin (right) plots showing dynamic expression of genes of interest in glial cells at Non-DEN, DEN2d, DEN5d, and DEN15d respectively.

**Supplementary Figure 7.**
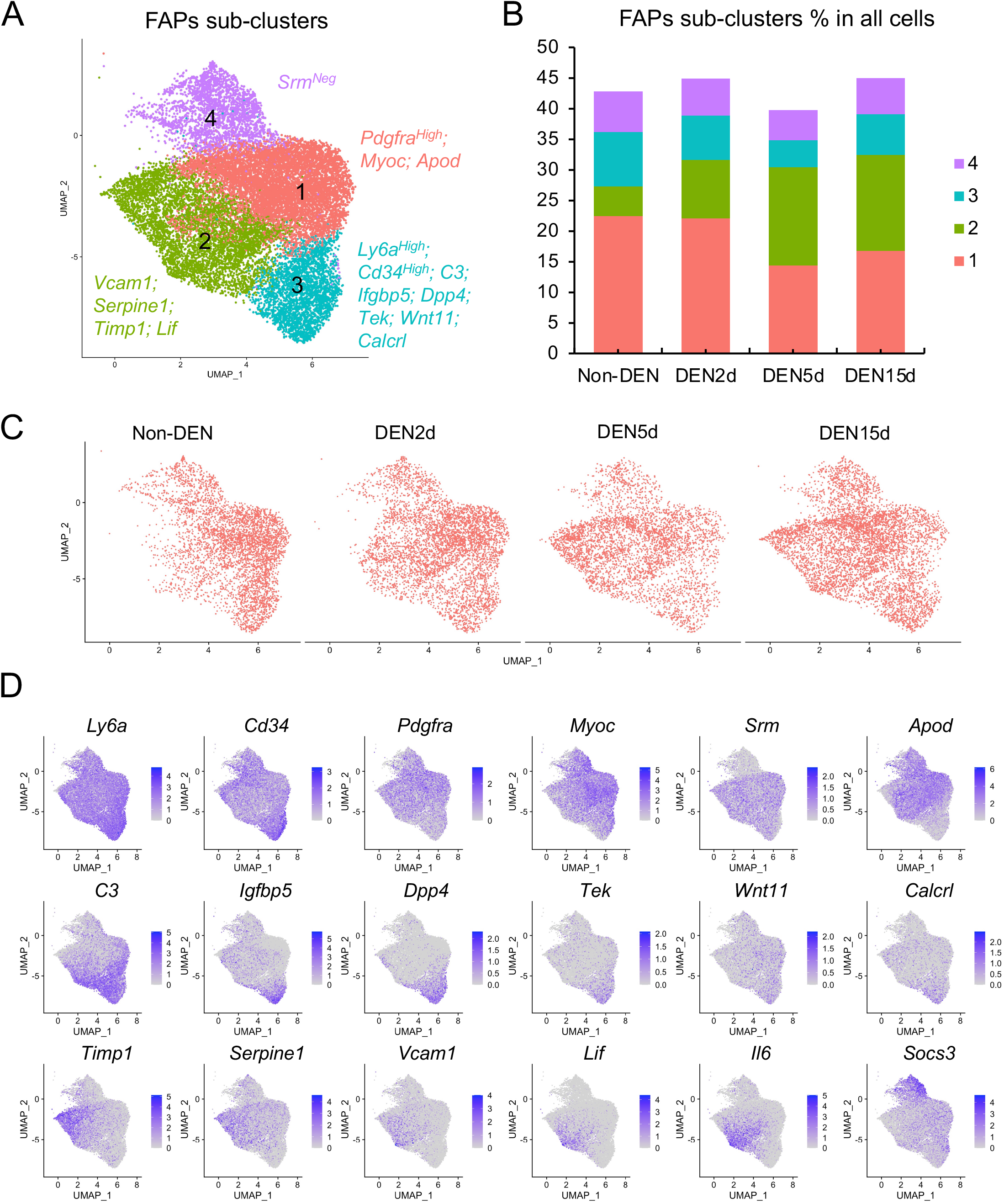
(A) UMAP of sub-clusters within FAPs. (B) FAPs sub-clusters percentage per condition. (C) UMAP embedding subset from the whole dataset of FAPs in time-lapse across denervation time points. (D) UMAP embedding of FAPs showing the expression of selected genes associated with adopogenic and fibrogenic lineages.

**Supplementary Figure 8.**
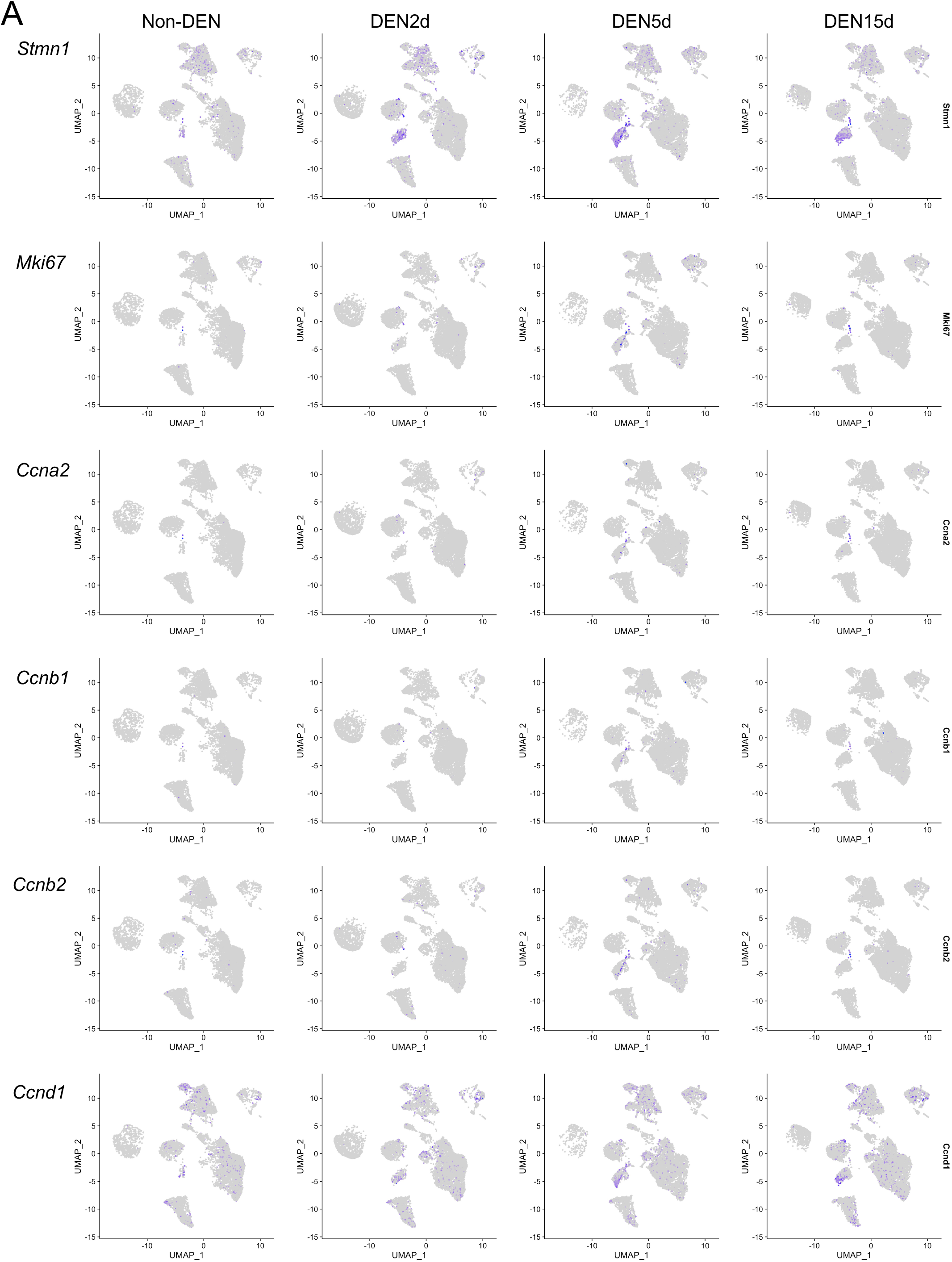
(A) UMAP embedding of all the single cells showing the expression of selected genes related to cell cycle.

**Supplementary Table 1.** Cell counts and percentages per condition for each cell population. Related to Figure 2A.

| Cell type | Non-DEN | DEN2d | DEN5d | DEN15d |
| --- | --- | --- | --- | --- |
| FAPs | 4502 (42.9%) | 4900 (45%) | 4124 (40.1%) | 5739 (45.1%) |
| Endothelial cells | 2115 (20.2%) | 1241 (11.4%) | 1927 (18.7%) | 1946 (15.3%) |
| Tenocytes | 1310 (12.5%) | 1188 (10.9%) | 1149 (11.2%) | 1029 (8.1%) |
| MuSCs | 764 (7.3%) | 858 (7.9%) | 703 (6.8%) | 901 (7.1%) |
| Inflammatory cells | 314 (3%) | 460 (4.2%) | 820 (8%) | 1042 (8.2%) |
| Myonuclei | 721 (6.9%) | 1223 (11.2%) | 95 (0.9%) | 46 (0.4%) |
| Glial cells | 93 (0.9%) | 241 (2.2%) | 593 (5.8%) | 683 (5.4%) |
| Myotenocytes | 239 (2.3%) | 227 (2.1%) | 223 (2.2%) | 722 (5.7%) |
| Activated Fibroblasts | 122 (1.2%) | 237 (2.2%) | 361 (3.5%) | 276 (2.2%) |
| Pericytes | 187 (1.8%) | 127 (1.2%) | 142 (1.4%) | 220 (1.7%) |
| SMMCs | 129 (1.2%) | 178 (1.6%) | 148 (1.4%) | 109 (0.9%) |
| **Total** | **10,496** | **10,880** | **10,285** | **12,713** |

**Supplementary Table 2.**
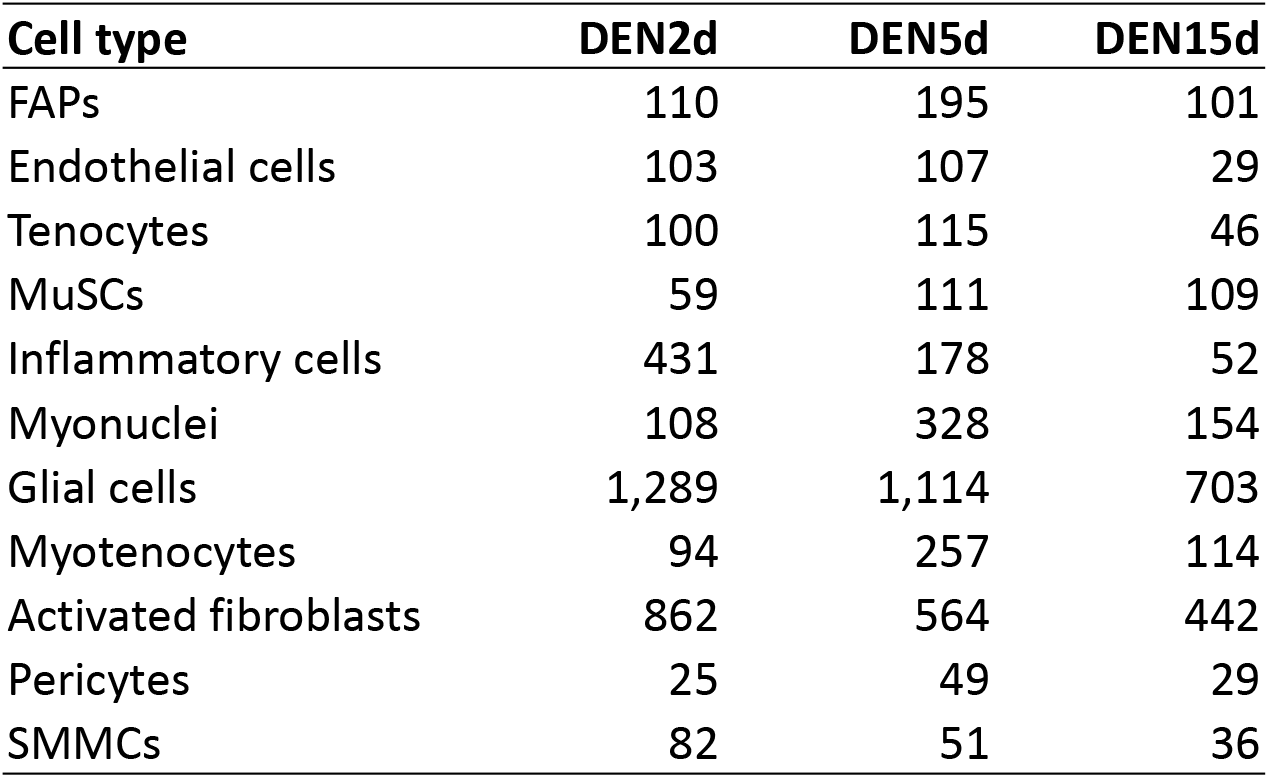
DE genes per condition for each cell population. Related to Figure 2B.

**Supplementary Table 3.**
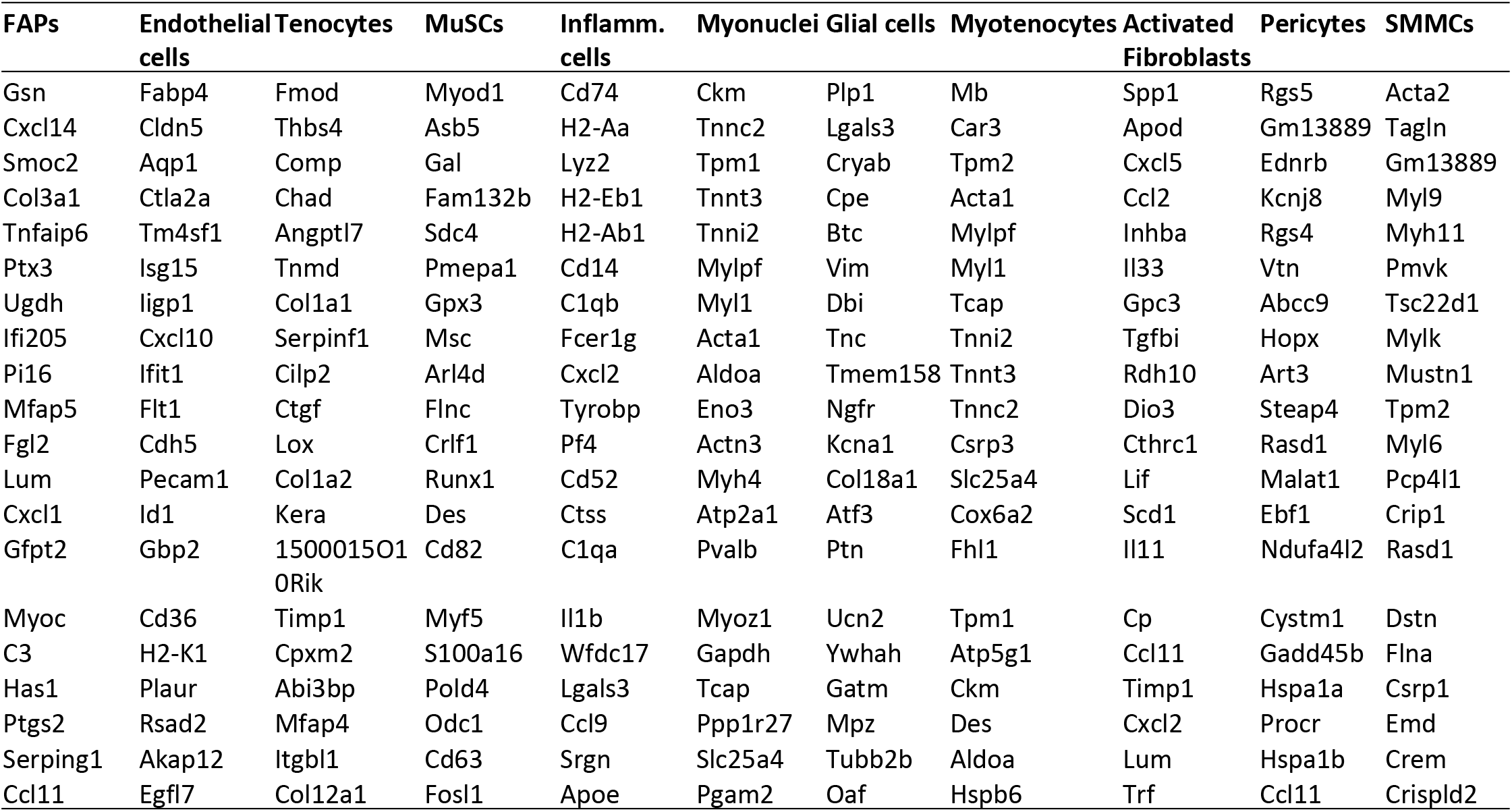
Top 20 marker genes for each cell population.

**Supplementary Table 4.** IPA comparison of DE genes (P-value-adjusted < 0.01) in glial cells at DEN2d, DEN5d, and DEN15d respectively compared with Non-DEN. Pathways significantly enriched in either time point with absolute z score > 1 were shown.

**Supplementary Table 5.** List of common differentially expressed (DE) genes in high Stmn1 expressing cells, within and between MuSCs and muscle-glial derived cells, across all conditions. High Stmn1 expressing cells were defined by normalized count >2, as compared to Stmn1 dim counterparts in MuSCs and glial cells.

| Gene |  |  |  |  |
| --- | --- | --- | --- | --- |
| Stmn1 | Atxn10 | Irf2bpl | Hmgn2 | Hspa5 |
| Lgals1 | Ppia | Gpc1 | Nes | Fgl2 |
| Fabp5 | Ppp1ca | Cdk4 | Prdx4 | Cebpb |
| Ccnd1 | Tmem256 | Sh3bgrl3 | Prc1 | Srsf2 |
| Gltp | Ndufa4 | 2700060E02Rik | Hist1h1c | Zbtb20 |
| Bok | 2700094K13Rik | Fscn1 | Arl6ip1 | Gsn |
| Cav2 | Dag1 | Dap | Tuba1a | Top1 |
| Mapk3 | Npc2 | Anapc11 | Tuba1b | Junb |
| Taldo1 | S100a4 | Angptl4 | H2afz | Tra2a |
| Vim | Atpif1 | Plp2 | Ifitm3 | Ccnl1 |
| Lgals3 | Mxra7 | S100a16 | Dcn | Clic4 |
| Prdx2 | Akr1a1 | Phlda3 | Malat1 | Ugp2 |
| Tmsb10 | Ndufa13 | Clic1 | H2-D1 | Jund |
| Nt5dc2 | Coro1c | Plec | Mt2 | Btg1 |
| Aes | Atp5o | Pcbp4 | Igfbp7 | Id3 |
| Tpm1 | Sdhc | Csrp2 | H2-K1 | Apoe |
| Hmgn3 | Ndufb8 | Pkm | Ifrd1 | Ppp1r15a |
| Ndufs2 | Commd1 | Sptssa | Dusp1 | Sdc4 |
| Calm3 | Tubb5 | Itga7 | Uap1 | Pdlim4 |

**Supplementary Table 6.** IPA comparison of DE genes (P-value-adjusted < 0.01) in inflammatory cells at DEN2d, DEN5d, and DEN15d respectively compared with Non-DEN. Pathways significantly enriched in either time point with absolute z score > 1 were shown.

